# In Silico Proteomics Approach Towards the Identification of Potential Novel Drug Targets Against *Cryptococcus gattii*

**DOI:** 10.1101/2022.07.29.502060

**Authors:** Tanjin Barketullah Robin, Nurul Amin Rani, Nadim Ahmed, Anindita Ash Prome, Md. Nazmul Islam Bappy, Foeaz Ahmed

**Affiliations:** Faculty of Biotechnology and Genetic Engineering, Sylhet Agricultural University, Sylhet-3100; Department of Molecular Biology and Genetic Engineering, Sylhet Agricultural University, Sylhet-3100

**Keywords:** *Cryptococcus gattii*, In Silico, ADME, Molecular docking, Paralogous, Drug targets, PBMCs

## Abstract

Cryptococcosis is a condition caused by inhaling *Cryptococcus gattii*, the tiny fungus from the environment. It is thought that the pathogen C. *gattii* is clinically more virulent than C. *neoformans* and could be a vicious agent in coming decades. It can enter the host’s brain and harm human peripheral blood mononuclear cells’ DNA (PBMCs). It is vital to investigate potential alternative medications to treat this disease since global antifungal resistance preventing Cryptococci infections is on the rise, leading to treatment failure. In order to find effective novel drug targets for C. *gattii*, a comprehensive novel approach has been used in conjunction with in silico analysis. Among 6561 proteins of C. *gattii* we have found three druggable proteins (XP 003194316.1, XP 003197297.1, XP 003197520.1) after completing a series of steps including exclusion of paralogs, human homologs, non-essential and human microbiome homologs proteins. These three proteins are involved in pathogen specific pathways, and can be targeted for drugs to eliminate the pathogen from the host. The subcellular locations and their interactions with a high number of proteins also demonstrate their eligibility as potential drug targets. We have approached their secondary, tertiary model and docked them with 21 potential antifungal plant metabolites. From the molecular docking analysis, we found Amentoflavone, Baicalin, Rutin and Viniferin to be the most effective drugs to stop such proteins because of their increased binding affinity. Correspondingly, the drugs showed proper ADME properties and also analyzed to be safe (Figure 9, Table 6). Moreover, these potential drugs can successfully be used in the treatment of Cryptococcosis caused by the fungus *Cryptococcus gattii*. In vivo trail is highly recommended for further prospection.

## Introduction

*Cryptococcus gattii* is a basidiomycetous yeast under the class Tremellomycetes and family Cryptococcaceae. It is an asexual budding yeast found in the environment and within human and animal hosts (Fraser et al., 2005). Sexual reproduction is possible in cells with the same mating type as well as cells with the opposite mating type. Yeast cells go through a dimorphic transition to hyphal growth, producing a mycelium and basidiospores, during sexual development. It is still unknown if the disease-causing propagules of C. *gattii* are sexually produced basidiospores, dehydrated yeast cells, or sexually produced blastospores. C. *gattii* is predominantly found in tropical and subtropical parts of the world, where it lives in soil and on certain trees.(Springer & Chaturvedi, 2010) Genomes of this organism range in size from 16 to 19 Mb, contain 14 chromosomes, and have a bipolar mating system (Montoya et al., 2021). This tiny fungus enters the body through inhalation from the environment, cause infection to lung and nervous system. The infection is called cryptococcosis. Though Cryptococcus neoformans also cause cryptococcosis, but infection of C. *gattii* is more dangerous as it is capable of infecting healthy individuals. C. *gattii* is more often associated with pulmonary cryptococcosis and occurs in immunologically competent individuals (Abee et al., 2012). Cryptococcal meningitis is the medical term for brain infections brought on by Cryptococcus *gattii* which frequently results in fatal consequences. The endemic pathogen Cryptococcus *gattii* infection started in the 1990s in British Columbia, Canada, and then spread to the Pacific Northwest of the United States and exists in Australia too. The recent Pacific Northwest outbreak of C. *gattii* infection serves as a reminder that it is a significant emerging fungal pathogen with the ability to adapt to new potential environmental conditions. C. *gattii* isolates discovered on Vancouver Island seem to be hypervirulent, as the number of affected individuals each year between 2002 and 2005 was more than 36 times greater on Vancouver Island compared to endemic regions such as Australia (Kidd et al., 2004; MacDougall et al., 2007). Moreover, the recent major advances in molecular technology have enhanced our understanding of ecology, epidemiology, and clinical associations of C. *gattii* as well as the whole genomic information.

Common drugs used in treatment of *Cryptococcus gattii* is Amphotericin B alone or in combination with flucytosine, as well as fluconazole alone or in combination with flucytosine, have both been used in the treatment of cryptococcosis in humans and nonhuman primates (Lowenstine & Osborn, 2012). Amphotericin B, either in its conventional or lipid formulation, is administered by extended intravenous therapy for a minimum of 6-8 weeks. Flucytosine used orally or intravenously enhances response rates. For at least six months, people with infections must take prescription antifungal medications; the kind of therapy relies typically on the severity of the illness and the areas of the body that are affected. Fluconazole is often used to treat asymptomatic infections and mild to severe lung infections. Fluconazole resistance in Cryptococcus has been steadily rising, from 7.3 percent from 1997 to 2000 to 10.9 percent from 2001 to 2004 and 11.7 percent from 2005 to 2007, according to a surveillance study (Mpoza et al., 2018). It is alarming that resistance is developing quickly against these medications. So we need new drug that can work more effectively than fluconazole. Drug discovery has benefited from developments in computational biology and bioinformatics methods using omics data such as proteomics, metabolomics, and genomics since they cut the cost and time required for in vivo and wet-lab drug design screening (Lin et al., 2020). Subtractive genomics is one of these bioinformatics methods, which compares the proteomes of the host and the pathogen to find non-host proteins with particular metabolic pathways that are essential to the survival of the pathogens. These essential proteins that displays no cross-reactivity with the host has been proposed as potential drug targets (Hasan et al., 2020). In this study we are going to identify the potential drug targets and suggest effective drug that can take up arms against *Cryptococcus gattii* using computational biology.

## Material and Methods

The unique drug targets of *Cryptococcus gattii* and prospective medications to prevent those distinct targets were examined using a variety of web servers and bioinformatics tools.

### 2.1 Retrieving Whole Proteomics of *Cryptococcus gattii*

The whole proteome of *Cryptococcus gattii* (assembly ASM18594v1) was obtained through the National Center for Biotechnology Information (https://www.ncbi.nlm.nih.gov/genome) server.

### 2.2 Exclusion of Duplicate and Mini Proteins

The extracted proteome of C. *gattii* was uploaded to the CD-HIT web-server (http://weizhong-lab.ucsd.edu/cdhitsuite/cgi-bin/index.cgi?cmd=cd-hit), in order to detect duplicate proteins. The “sequence identity cut-off” was fixed to 60 % (0.6) for the removal of redundant sequences of protein. Protein sequences having fewer than hundred amino acid are vital for a variety of regulative processes and other biological functions; therefor to omit those the “length of sequence to skip” was fixed to hundred (Gupta et al., 1948). While amino acids with larger sequences are usually found to evolve in crucial metabolic functions (Haag et al., 2012).

### 2.3 Detection of Human Proteome Non-Homologous Protein Sequences

Avoiding functional resemblance to the human proteome was the focus of this step, the idea is to avoid the bindings of drug with the host homologous proteins’ active sites. The Ensemble server was utilized to run BLASTp opposed to the human (Homo sapiens) Refseq proteome, non-paralogous pathogen proteins were examined (Zerbino et al., 2018). The proteins were deemed to be homologous to the host when any consequential hits greater than the threshold level 10^−4^ were discovered. The objective here is to eliminate any possibilities of resemblance with human proteome.

### 2.4 Detection of essential unique proteins

The Database of Essential Genes (DEG) was used to analyze the proteins that are not homologous with host. (Luo et al., 2014). The DEG 15.2 server’s entire collection of organism strains were chosen, and BLASTp was run using the parameters of a threshold level of 10 and a bit score of at least 100. Proteins that were hit with an identity of 25 percent or less and an expectation value of 10 to 100 were kept because they were thought to be crucial for the pathogen. A bacterium’s minimal genome is made up of the byproducts of its essential genes, which are important in the field of synthetic biology (Kanehisa & Goto, 2000).

### 2.5 Identification of Unique Metabolic Pathway

The metabolic pathways for both *Cryptococcus gattii* and humans were acquired by utilizing the KEGG PATHWAY server (https://www.genome.jp/kegg/pathway.html) through inputting the appropriate 3 letter organism codes at KEGG which are ‘cgi’ and ‘hsa’ respectively for *Cryptococcus gattii* and H. *sapiens* (Ogata et al., 1999). Then, manual comparison was used to screen the bacterium’s unique pathways. The KAAS server(https://www.genome.jp/kaas-bin/kaas_main) of KEEG database’s and BLASTp was applied to identify the KO number of crucial host non-homologous proteins. The sequences that are merely involved in the organism’s distinct metabolic pathways were identified using the KO number by utilizing the KO server (https://www.genome.jp/kegg/ko.html) (Damte et al., 2013). Protein sequences having non-similar pathways were retrieved for the further study.

### 2.6 Determining Host Microbiome Non-similar Proteins

A vast amount of helpful microorganisms presents in the host body that works to protect the body from unknown objects. Finding out whether the distinct metabolic pathway sequences of *Cryptococcus gattii* match the protein of these helpful microorganisms present in the host body is the objective of the host microbiome non-similar protein study. So, NCBI BLAST server is utilized to BLAST the distinct metabolic pathway protein sequences opposing Bioproject-43021 while having the cut of score at 0.005 (Turnbaugh et al., 2007). For succeeding phases, sequences that share fewer than 45% of their similarities were retained.

### 2.7 Identifying Novel Therapeutic Targets

To emphasize the distinctiveness of the chosen protein sequences as pharmacological targets, sequences from the earlier stages were examined at the Drug Bank server database (https://www.drugbank.ca/structures/search/bonds/sequence) (Wishart et al., 2018). Targets show how draggable they are, while the absence of targets shows how distinctive the proteins classified as “new targets” are (Knox et al., 2011). Throughout the action, the predefined settings for each variable were kept as the same.

### 2.8 Cellular Localization

Although cytoplasmic proteins have the potential to be therapeutic targets, proteins in the membrane can be employed as drug targets as well as vaccine contender (Mahmud et al., 2019). To anticipate the subcellular localization of the distinct therapeutic targets, the CELLO v.2.5: subcellular localization predictor server (http://cello.life.nctu.edu.tw) was employed.

### 2.9 Protein-Protein Interactome Analysis

To analyze the interaction network of the proteins of distinct drug targets the STRING 11.5 software (https://string-db.org) server was utilized (Szklarczyk et al., 2019). High confidence interactors with scores under 0.700 are present in the protein network. to remove false positive and negative results. When the query protein is removed, the amount of edges and nodes which means interconnections and interconnecting proteins will show how it affects the organism’s biological process (Kushwaha & Shakya, 2010).

### 2.10 Predicting Primary, Tertiary and Protein’s Binding Sites

The chosen proteins’ structures were not available on the Protein Data Bank (RCSB PDB). In order to anticipate molecular modellings of distinct protein sequences, the I-TASSER server was utilized (Roy et al., 2010). GalaxyWEB server(http://galaxy.seoklab.org/cgi-bin/submit.cgi?type=REFINE) was then employed for refining these models (Ko et al., 2012). To find the best model, the Errat value as well as Ramachandran plot of each models were examined using Saves v6.0 server (https://saves.mbi.ucla.edu/) (Colovos & Yeates, 1993; Laskowski et al., 1996)

### 2.11 Assemble of Plant Metabolites

A collection of 21 metabolites from different plants that was discovered having anti-fungal effects was compiled after reading the literature. According to Kim et al. (2016), We first obtained the three dimensional structure of the selected metabolite in SDS format using the PubChem database (https://pubchem.ncbi.nlm.nih.gov), and utilized the Open Babel v2.3 program to transform them into Protein Data Bank (PDB) format.

### 2.12 Molecular Docking

In drug development, molecular docking can be used to forecast how small ligands will interact with large molecules (Kitchen et al., 2004). Docking was performed by utilizing the Server HDock (http://hdock.phys.hust.edu.cn/) because it allows for connection within therapeutic targets and potential treatments (Meng et al., 2011; Yan et al., 2020). The ligands were chemicals, and the receptors were proteins. Since fluconazole is currently employed to treat *Cryptococcus gattii*, the baseline metabolite for the docking analysis was considered as being it. Then, the binding sites of the metabolites were visualized and examined using the PyMOl v2.0 software (Wang et al., 2015).

### 2.13 Toxicity study and Pharmacoinformatic Analysis

The kinetics of drug exposure to tissue plays a major role in how a drug’s adsorption, distribution, metabolism, and excretion (ADME) characteristics seems related. Assessing ADME in the evaluation stage will lessen the likelihood of clinical failure due to pharmacokinetics. (Hay et al., 2014). The ADME characteristics of four leading metabolites were administered. using the SwissADME server (http://www.swissadme.ch/) (Daina & Zoete, 2016). Predictions were made by running the drugs after they had been uploaded to the server in SDF format and converted to SMILES. The BOILED-Egg model was further utilized to calculate how well the compounds studied penetrated the blood-brain barrier (BBB) (Daina & Zoete, 2016). Researchers utilized pkCSM, an online software that implies SMILES for the four leading metabolites, that were also sourced from the PubChem database, to anticipate the comparative negative effects of leading therapeutics. (Pires et al., 2015)

## Results

### 3.1 Exclusion of Paralog and Mini Proteins

We assembled the *Cryptococcus gattii* representative proteome (assembly ASM18594v1), that is consist of 6561 proteins, from the 8 genomes for *Cryptococcus gattii* that NCBI has available. After removing the paralogous sequences, the CD-HIT server found 6142 clusters at 60 percent identity, leaving 6142 large proteins that were not duplicates (Supplementary file S1)

### 3.2. Selecting host non-similar protein sequences

Protein sequences that are engaged in general cellular pathways for both bacterial and human had begun to resemble one another and arose as homolog (Hediger et al., 1989; Swango et al., 2000) As a result, drugs generated and used to interact with pathogen target proteins should prevent co-reactivity involving host homologous proteins. Ensemble genome platform 92 was utilized for operating BLASTp for all the 6142 proteins of C. *gattii* opposing to humans (*Homo sapiens*) proteins. The result was found to be 3800 proteins that displayed great hit more than threshold level along human proteome. A total of 3800 proteins showed significant hits with human proteins above the threshold. We removed these 3800 proteins assessing that those are human homologous, the other 2342 non-resemblance proteins were kept for further study (supplementary file S2).

### 3.3. Selecting Essential Protein Sequences

Antibacterial substances are frequently designed to interact with and obstruct vital gene products. Therefore, essential proteins are thought to be the most effective therapeutic targets (Zhang et al., 2004). Through dataset of essential proteins for bacteria that was maintained in the DEG 15.2 site, 216 proteins in total demonstrated significant hits. Assessing the proteins as vital that *Cryptococcus gattii* WM276 requires to survive, these 216 sequences were retrieved (supplementary file S3). Important genetic components that are unique to a microbe could serve as a therapeutic agent for that species specially. (Judson & Mekalanos, 2000)

### 3.4 Unique Metabolic Pathway Proteins Identification

There were 345 human metabolic pathways and 120 C. *gattii* metabolic pathways on the KEGG platform, with 17 metabolic pathways (Supplementary file S4) being unique to C. *gattii*. Therapeutic targets can be proteins involved in these distinct pathways. In the aftermath of a BLAST utilizing the KAAS database, it was found that 117 out of 216 essential non-homolog proteins were involved in both metabolic pathways and orthologs. These 117 proteins are essential for the proper functioning of metabolism, and 7 of them were found to be exclusively linked to specific pathways found only in C. *gattii*, mentioned in Table 1.

**Table 1:**
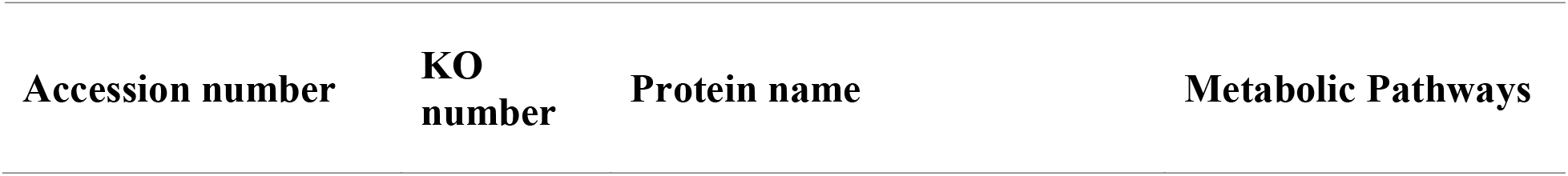
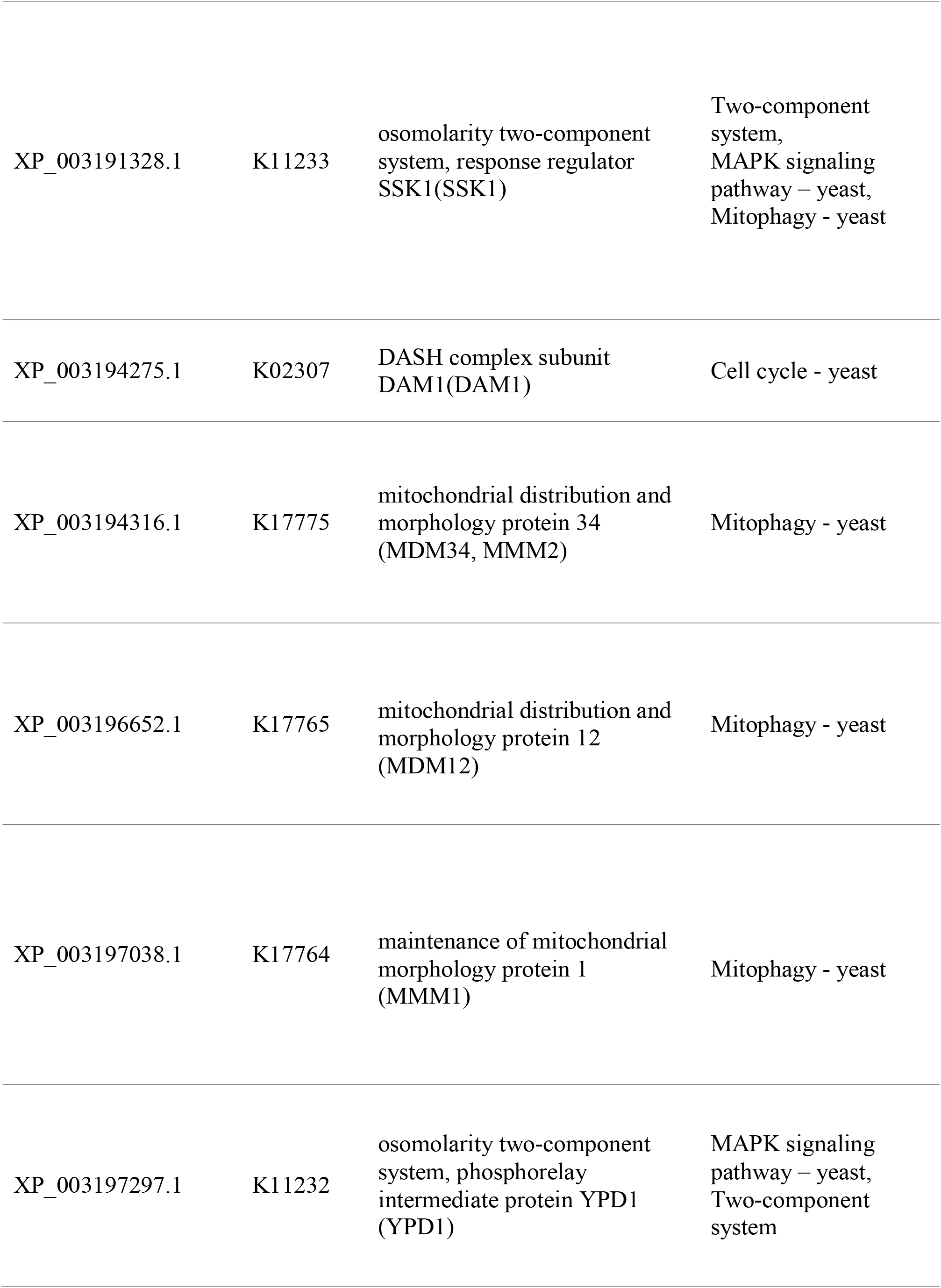
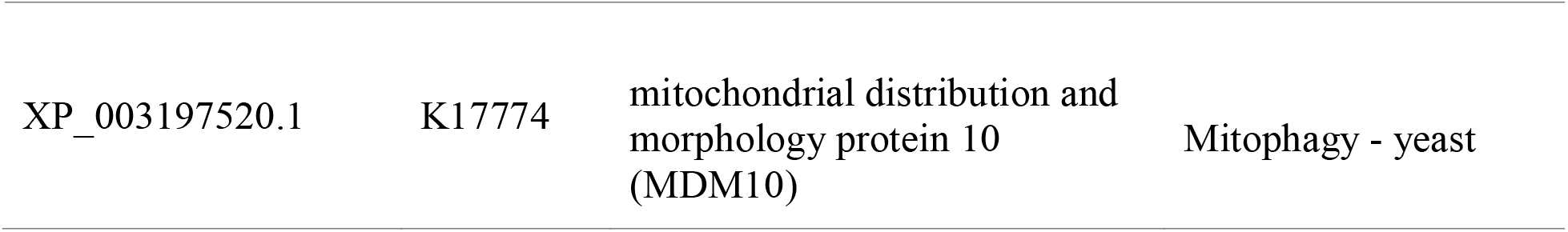
Proteins with Unique metabolic pathways and their KO number

| Accession number | KO number | Protein name | Metabolic Pathways |
| --- | --- | --- | --- |
| XP_003191328.1 | K11233 | osomolarity two-component system, response regulator SSK1(SSK1) | Two-component system, MAPK signaling pathway – yeast, Mitophagy - yeast |
| XP_003194275.1 | K02307 | DASH complex subunit DAM1(DAM1) | Cell cycle - yeast |
| XP_003194316.1 | K17775 | mitochondrial distribution and morphology protein 34 (MDM34, MMM2) | Mitophagy - yeast |
| XP_003196652.1 | K17765 | mitochondrial distribution and morphology protein 12 (MDM12) | Mitophagy - yeast |
| XP_003197038.1 | K17764 | maintenance of mitochondrial morphology protein 1 (MMM1) | Mitophagy - yeast |
| XP_003197297.1 | K11232 | osomolarity two-component system, phosphorelay intermediate protein YPD1 (YPD1) | MAPK signaling pathway – yeast, Two-component system |
| XP_003197520.1 | K17774 | mitochondrial distribution and morphology protein 10 (MDM10) | Mitophagy - yeast |

### 3.5 Drug Target Analysis and Host Microbiome Non-homology study

Six pathogen proteins were identified through microbiome analysis as having a resemblance of fewer to 45% with human microbial flora and having no similarity to authorized, conjectural and tested drug targets in the database of Drugbank. Three of which found to be extremely important for the metabolic cycle of C. *gattii*. They were proposed as the pathogen’s exclusive drug targets as a result (Table 2).

**Table 2:**
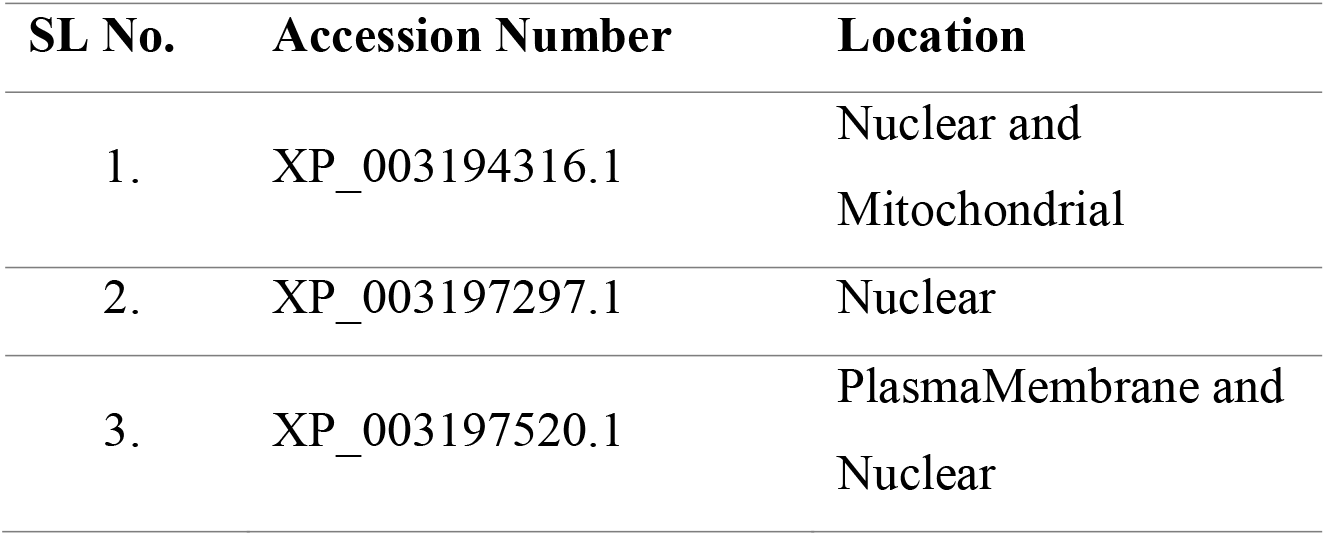
C. *gattii* specific drug targets and their localization.

### 3.6 Determining Subcellular Location

Either the nuclear mitochondria or the plasma membrane contains every one of the special pathway proteins. While XP 003194316.1 is found in the mitochondria, XP 003197297.1 is only present in nuclear regions. XP 003197520.1 is a protein found in the plasma membrane (Table 2). All of them were kept because they could be exploited as probable therapeutic agent in future research.

### 3.7 Protein-Protein Interactome Study

It is assumed that metabolically active proteins are promising therapeutic targets because of their engagement with other proteins (Cui et al., 2009). XP 003194316.1, XP 003197297.1, and XP 003197520.1 all demonstrate interconnections with Ten proteins, according to STRING as showed in Figure 2.

**Figure 01:**
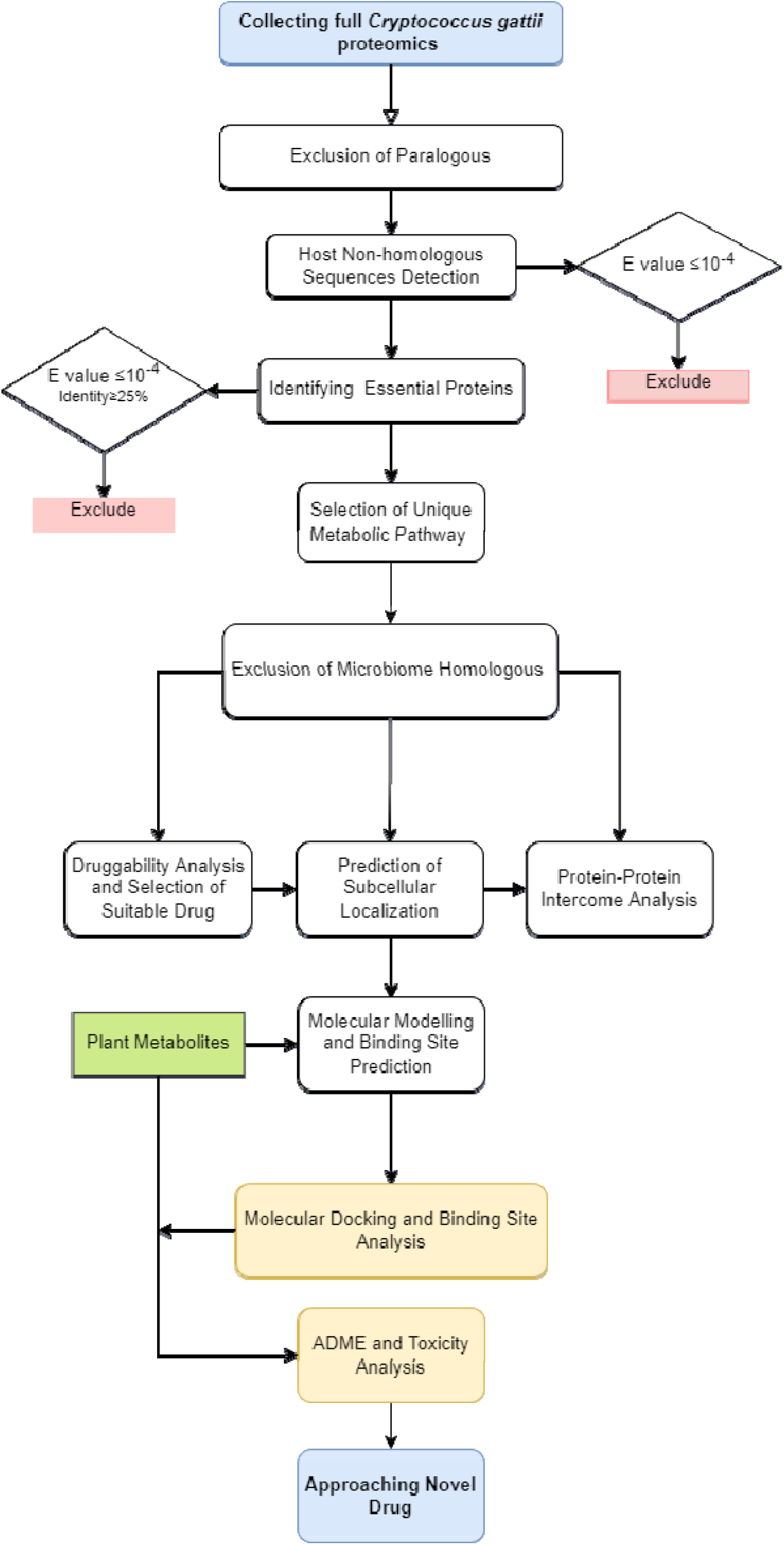
Schematic representation of the procedure

**Figure 2:**
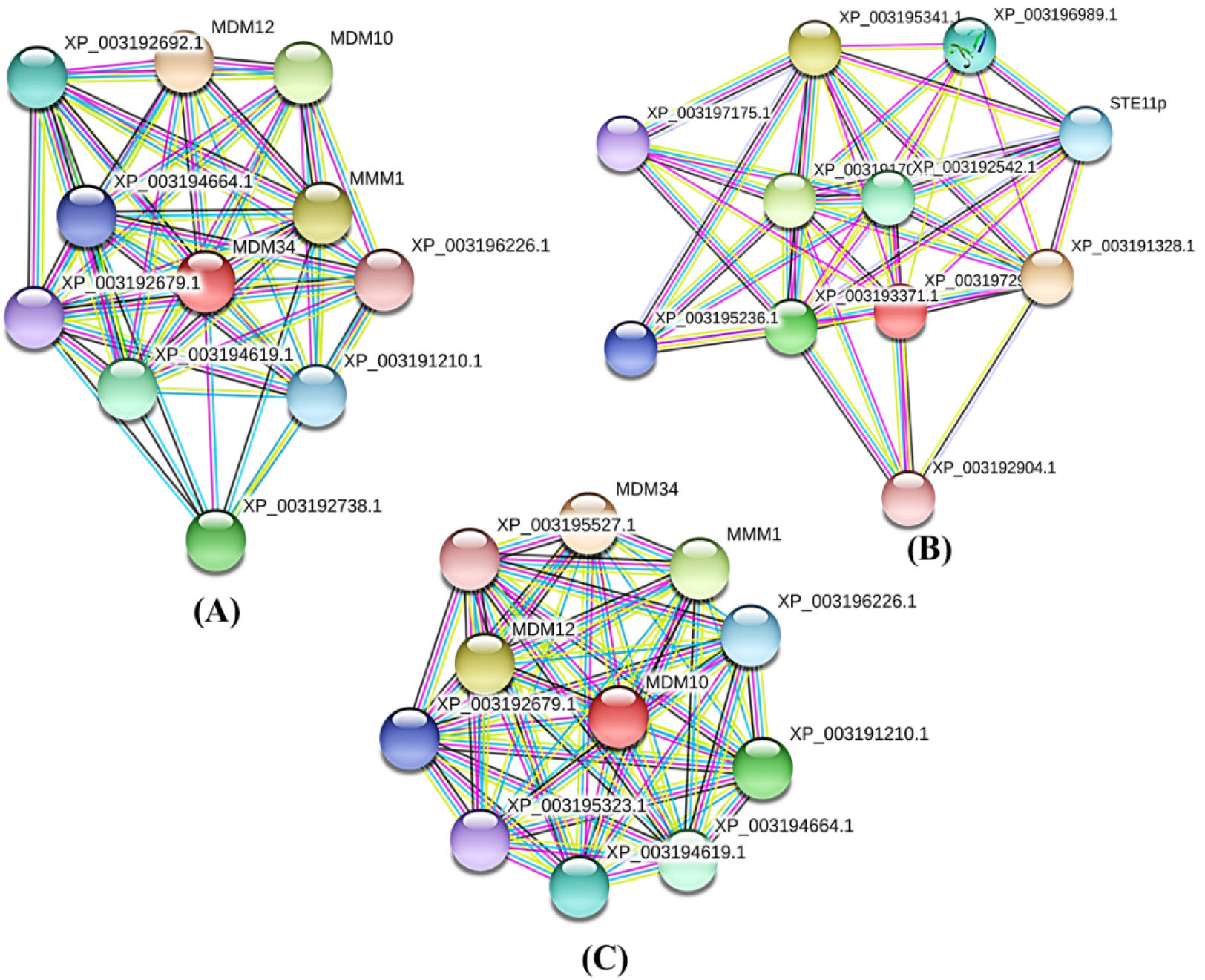
Interconnection of (A) XP 003194316.1, (B) XP 003197297.1, and (C) XP 003197520.1 with different proteins, the target protein is indicated in red color.

### 3.8 Molecular modelling Analysis

For each protein, 5 predicted patterns were offered by the I-TASSER server. Then By inspecting the ERRAT score and Ramachandran layout, the prime model was improved using the GalaxyWEB server. For each protein, this server provided 5 improved models. We eventually chose the best model by comparing their ERRAT grade level and Ramachandran layout assessment. Table 3, Figure 3, Figure 4, and Figure 5 provide descriptions of the outcome of the Ramachandran scheme and the ERRAT result.

**Table 3:** ERRAT values, Ramachandran plot results with residues for binding sites for refined protein model

| Accession ID | Errat value | Ramachandran plot result |  |
| --- | --- | --- | --- |
|  |  | Favored region | Disallowed region |
| XP 003194316.1 | 66.366 | 72.4% | 3.5% |
| XP 003197297.1 | 91.045 | 80.2% | 2.6% |
| XP 003197520.1 | 73.805 | 79.2% | 2.1% |

**Figure 3:**
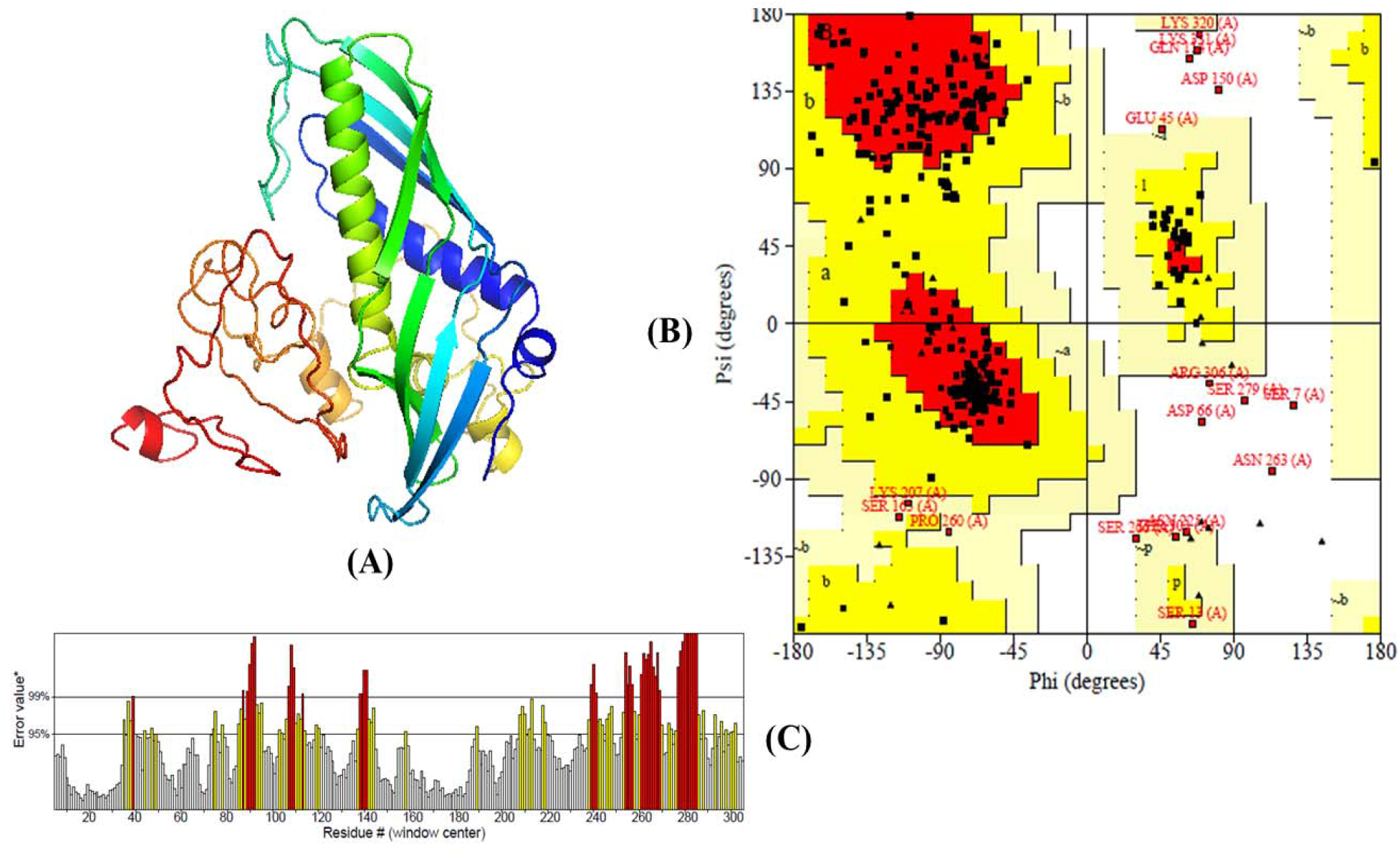
(A) Three-dimensional model, (B) Ramachandran layout and (C) The Errat value of XP 003194316.1

**Figure 4:**
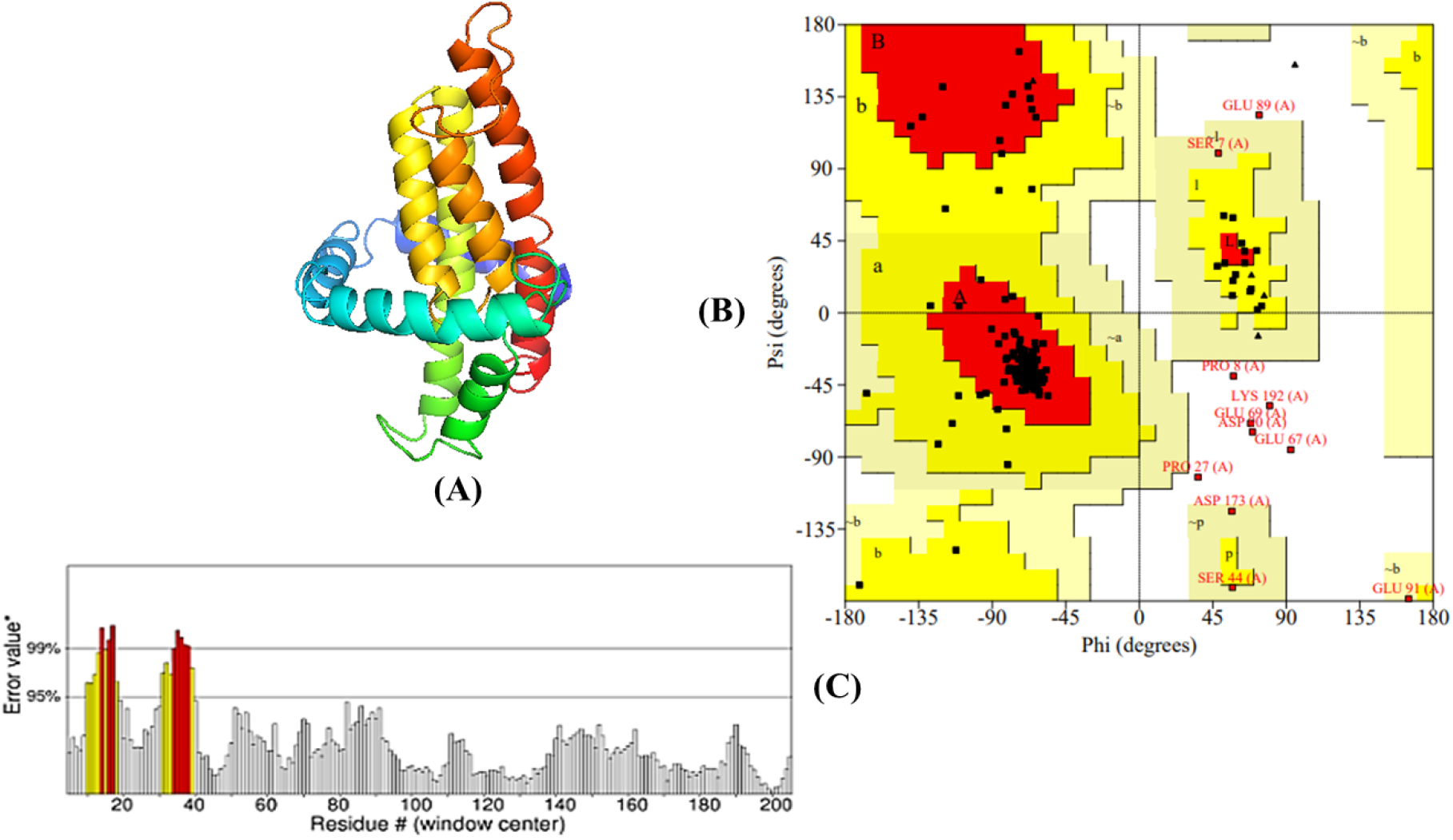
(A) Three-dimensional, (B) Ramachandran layout and (C) The Errat quality value of XP 003197297.1

**Figure 5:**
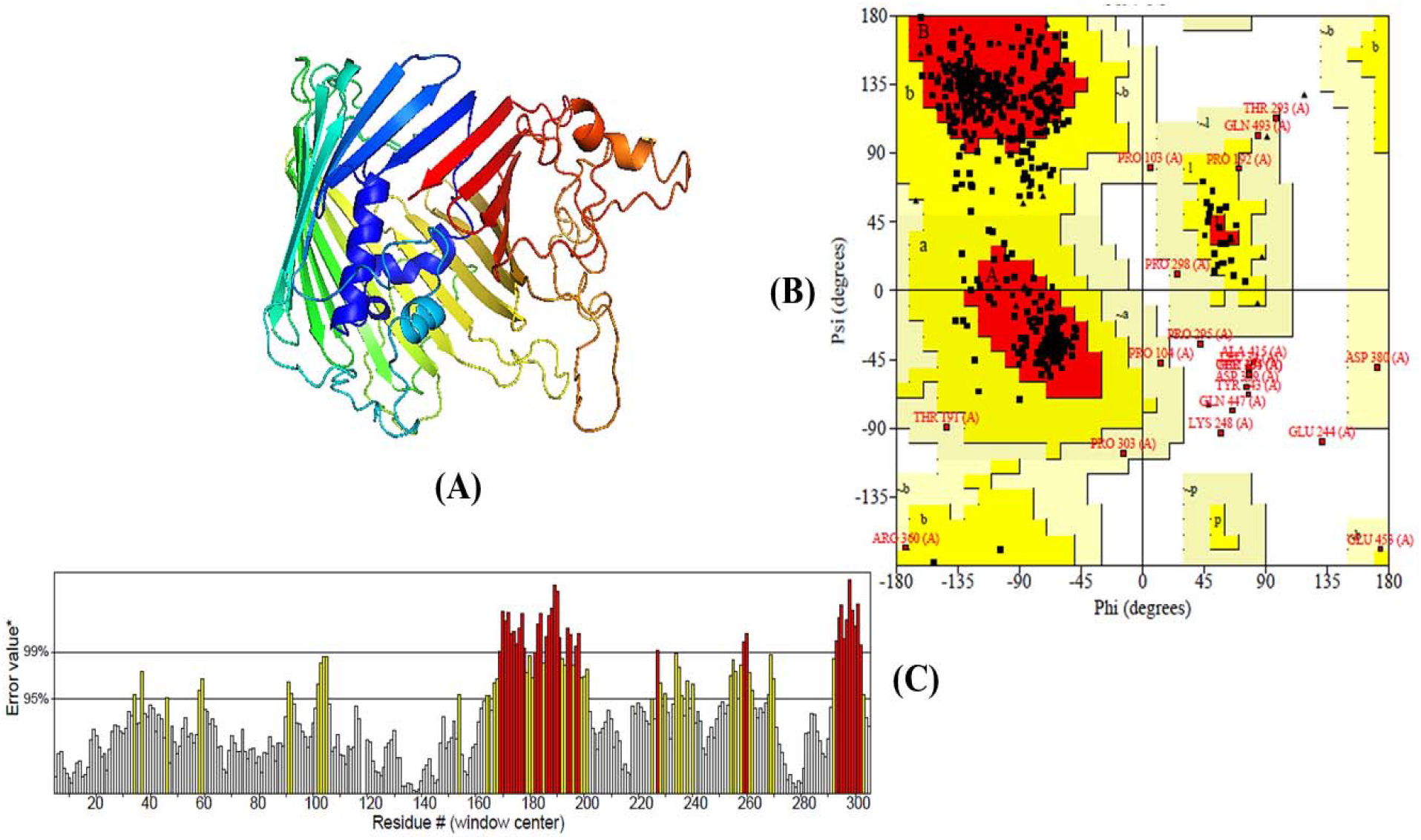
(A) Three-dimensional, (B) Ramachandran layout and (C) The Errat quality value of XP 003197520.1

### 3.9 Docking and Binding Site Assessment

Supplementary file S5 contains a list of the plant metabolites that have been collected. Every single one of our chosen special therapeutic targets docked opposed to every single metabolite (ligand) (Supplementary file S6). When docked with the chosen drug targets, Amentoflavone, Baicalin, Rutin, and Viniferin performed better than fluconazole. We therefore used the PyMol tool to further analyze the metabolites binding or interacting sites and listed the results in table 4, figure 6, figure 7, and figure 8.

**Table 4:**
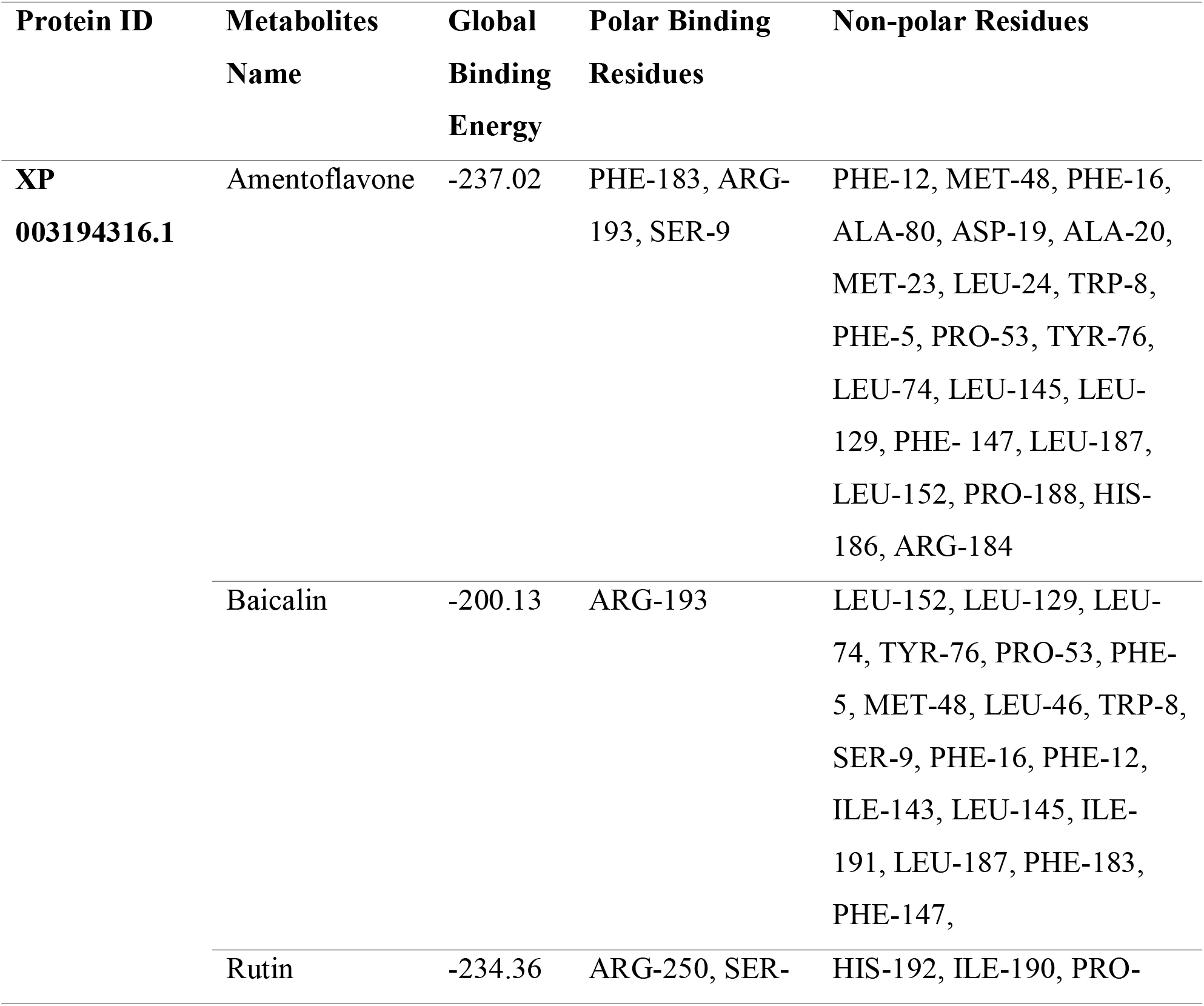
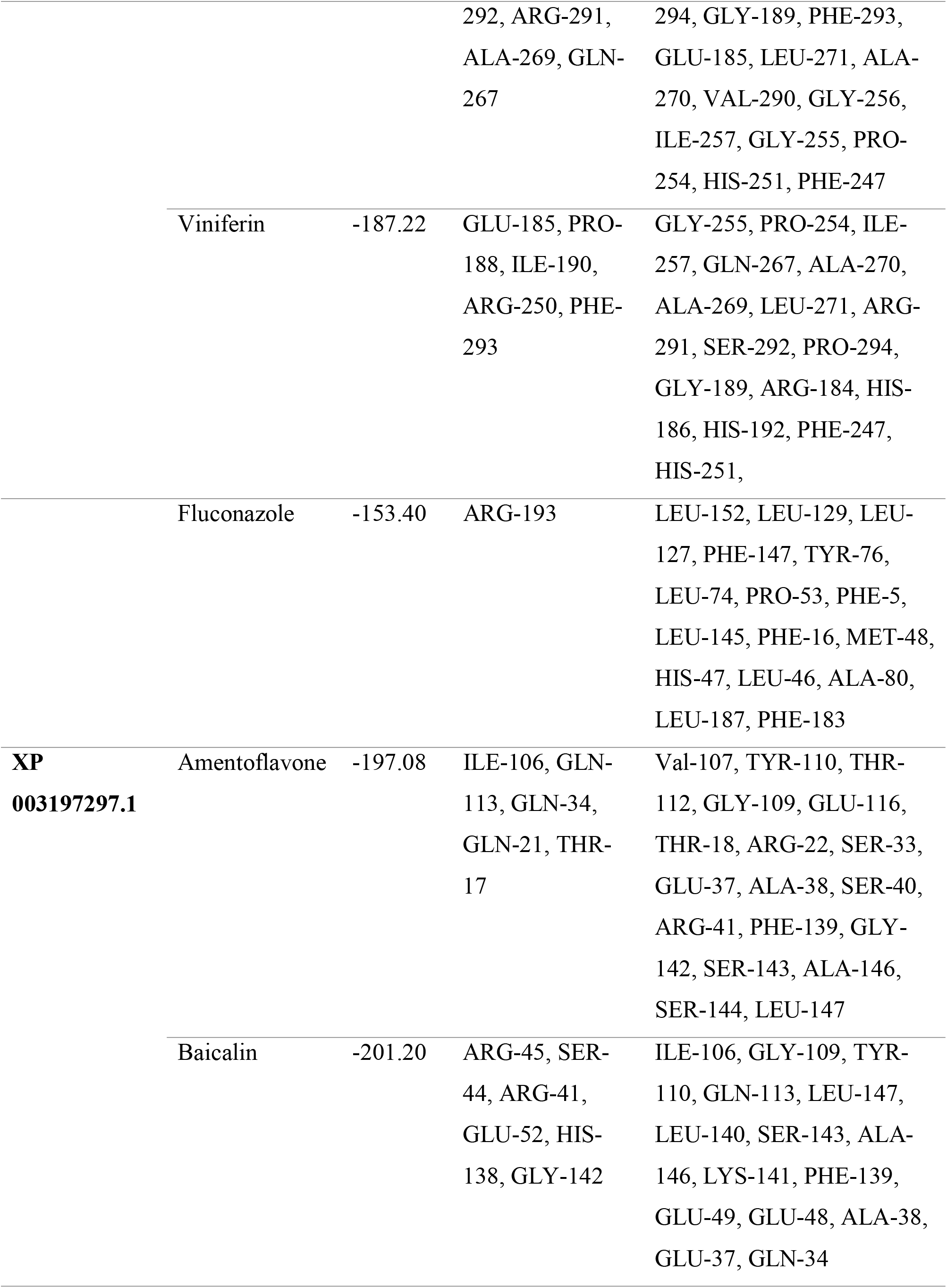
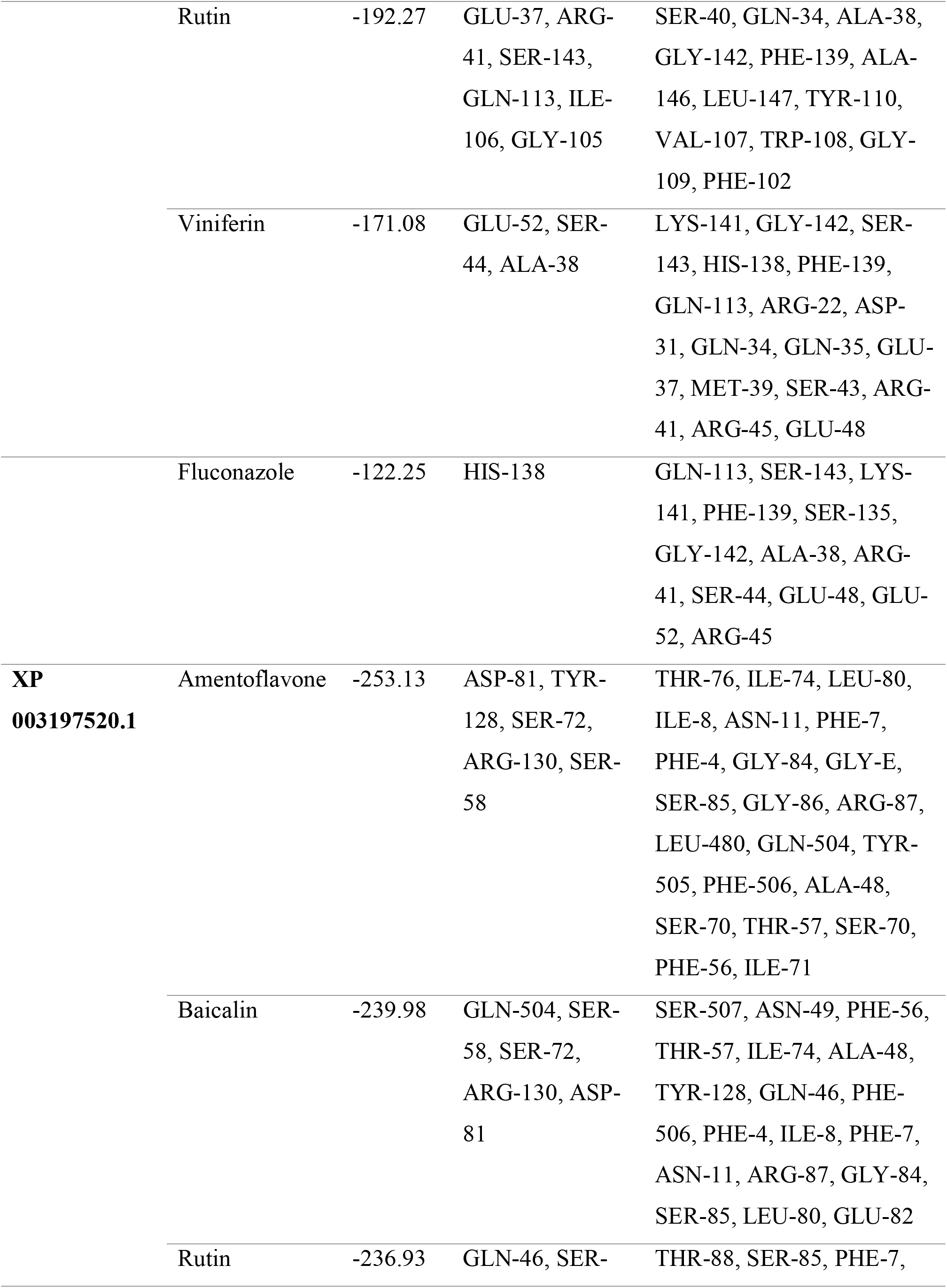
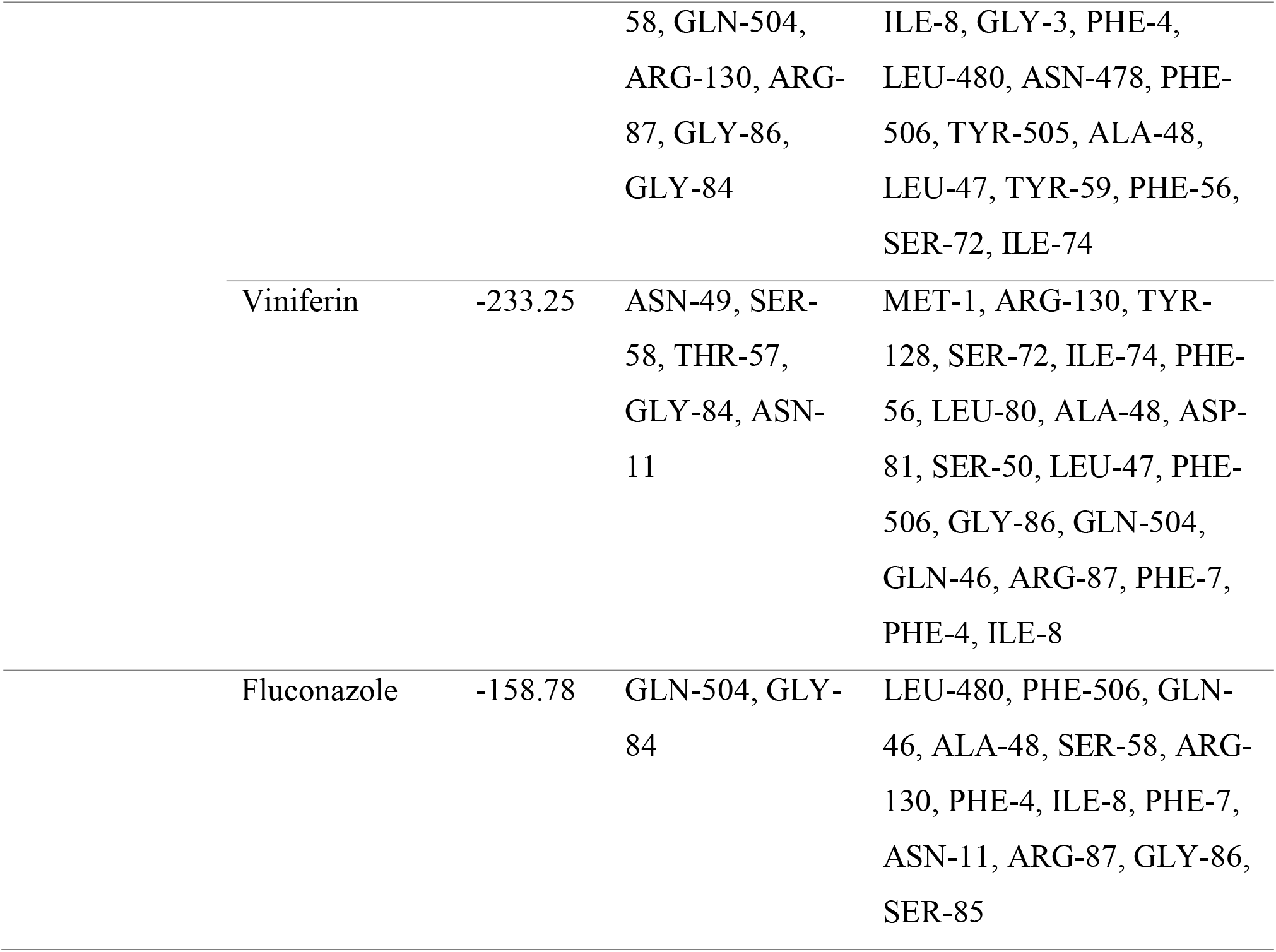
Binding regions of Amentoflavone, Baicalin, Rutin, and Viniferin

**Figure 6:**
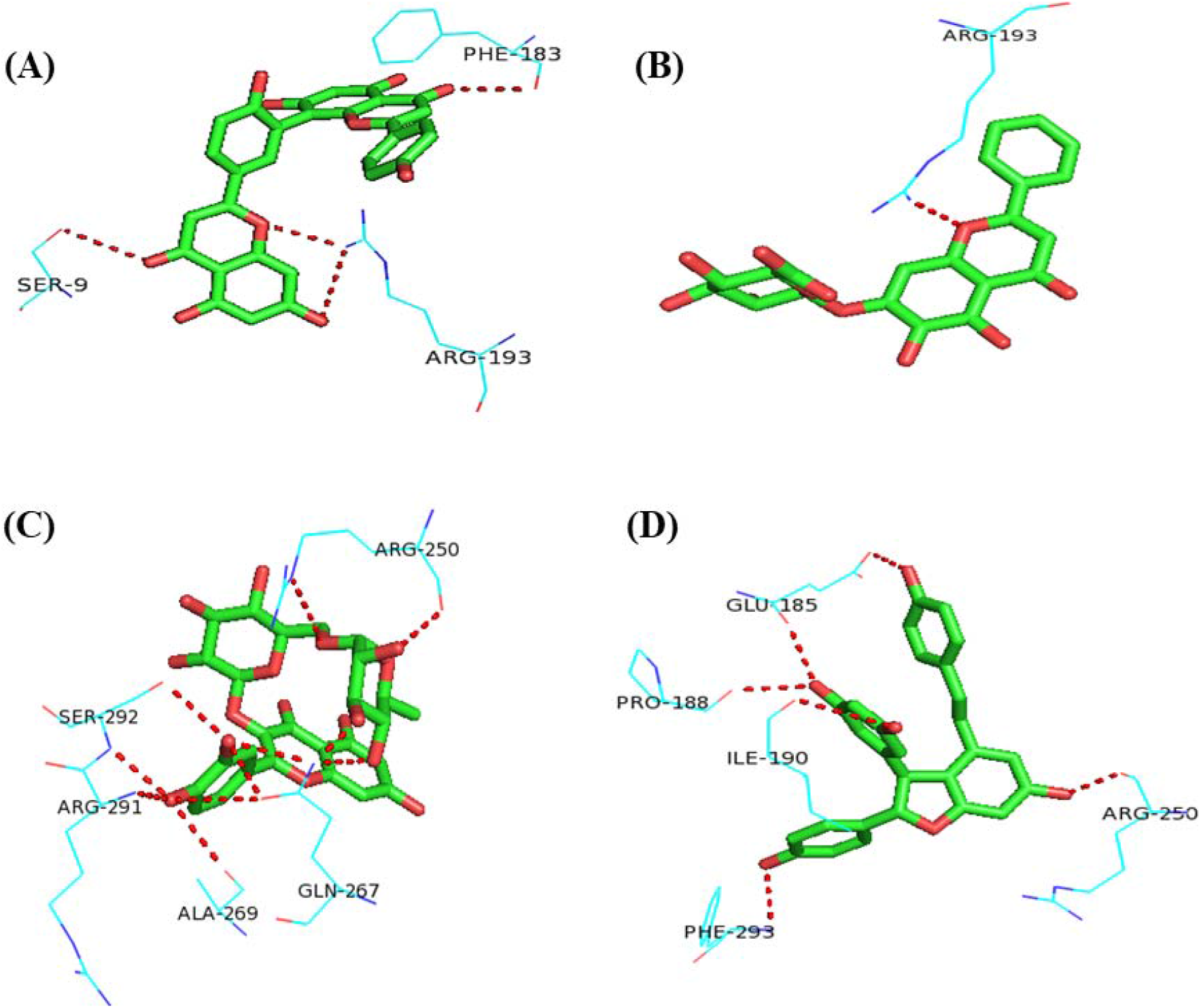
XP 003194316.1 with (A) Amentoflavone, (B) Baicalin, (C) Rutin, and (D) Viniferin and their polar binding residues.

**Figure 7:**
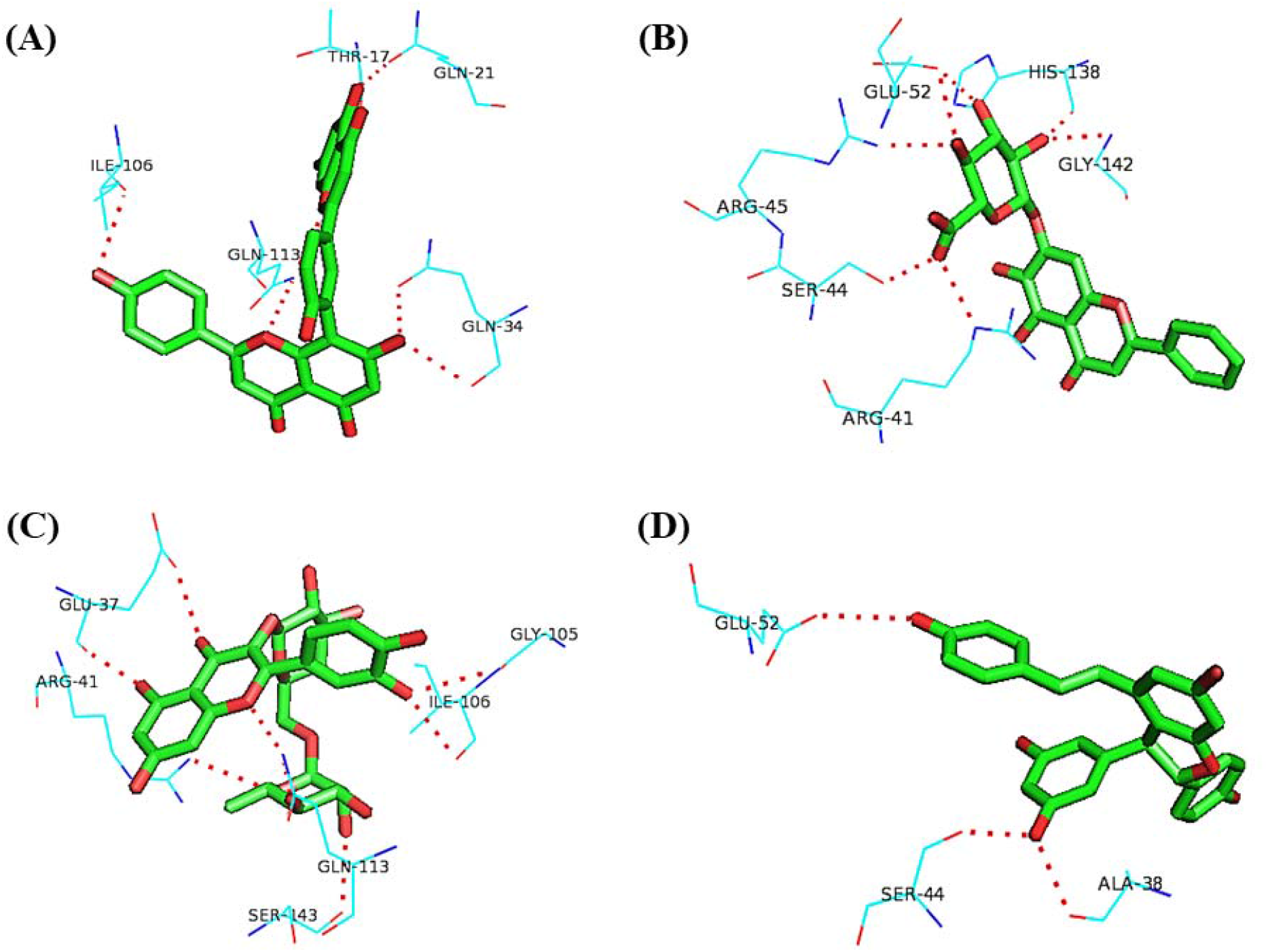
XP 003197297.1 with (A) Amentoflavone, (B) Baicalin, (C) Rutin, and (D) Viniferin their polar binding residues.

**Figure 8:**
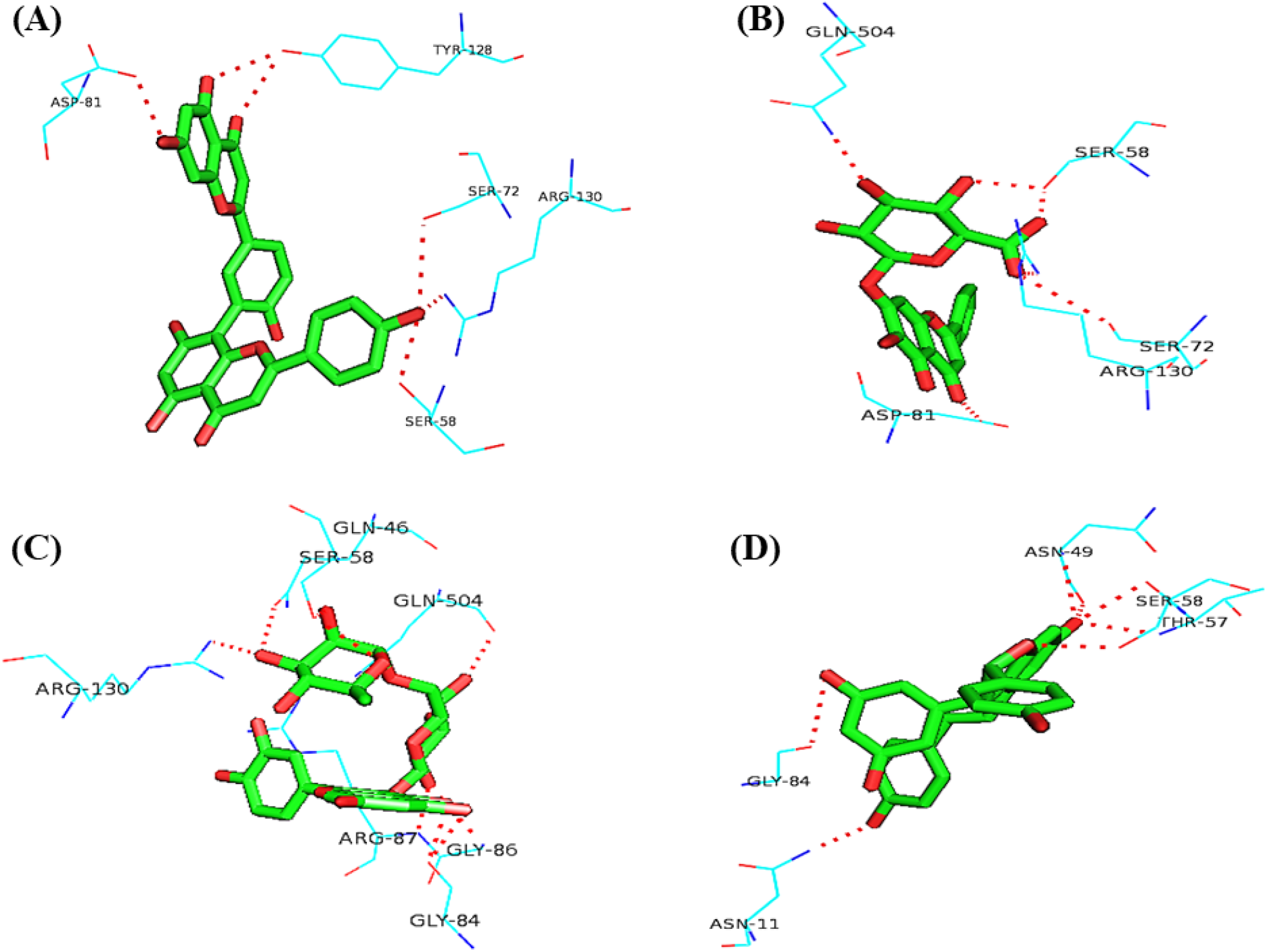
XP 003197520.1 with (A) Amentoflavone, (B) Baicalin, (C) Rutin, and (D) Viniferin their polar binding residues.

**Figure 9:**
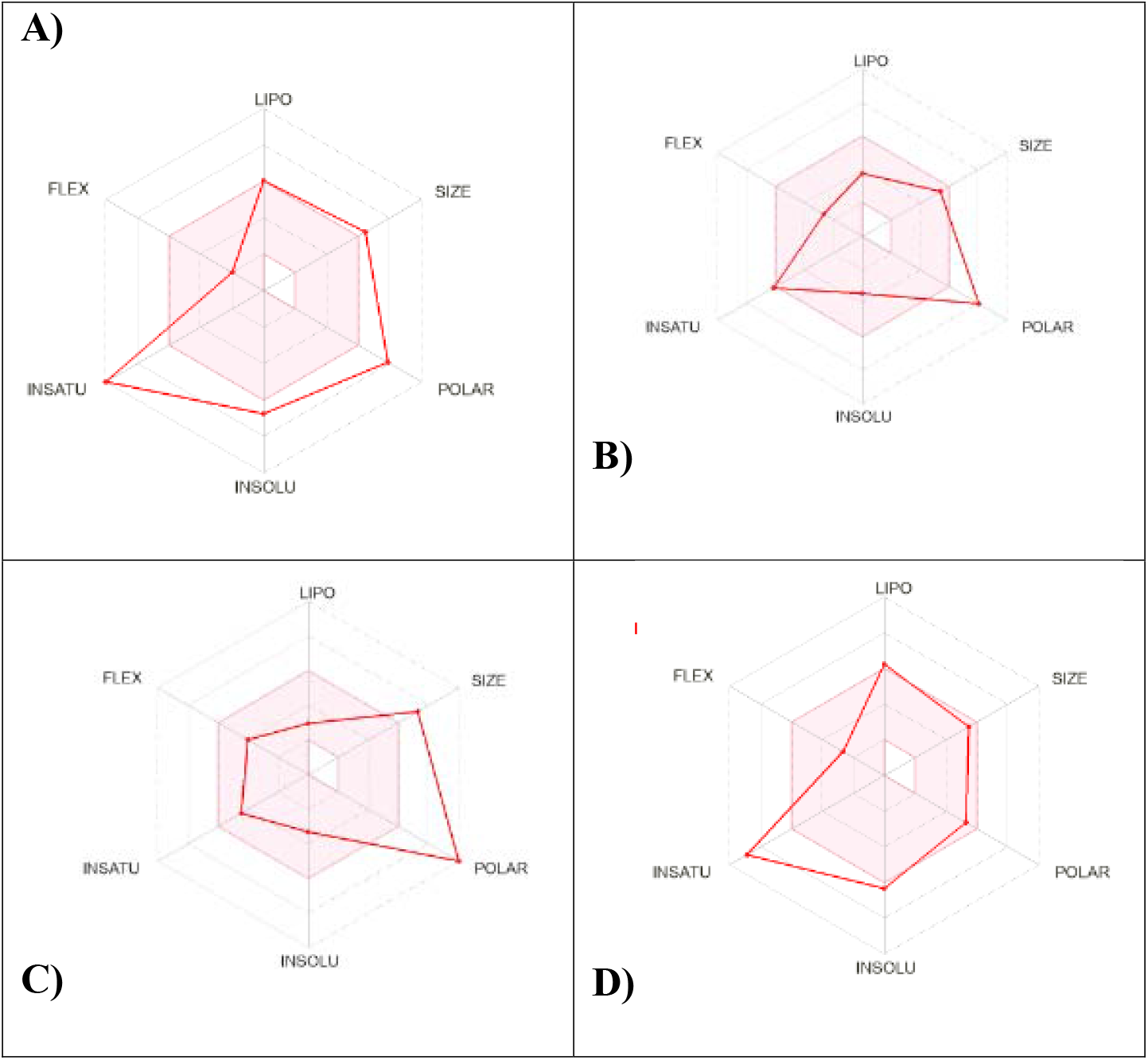
SwissADME properties of top metabolites

### 3.10 Pharmacoinformatic Analysis

Describing the therapeutic properties of the most effective antibiotics, the ADME characteristics of popular medications were evaluated (Table 5). In the digestive system, all of the metabolites were rapidly absorbed. These popular medications, Amentoflavone, Baicalin, Rutin, and Viniferin, did not contain any BBB. Aside from Viniferin, no medication was identified to inhibit CYP2C9. Additionally, Baicalin and rutin had good solubility while only Amentoflavone and viniferin displayed poor solubility, mentioned in Table 5

**Table 5:**
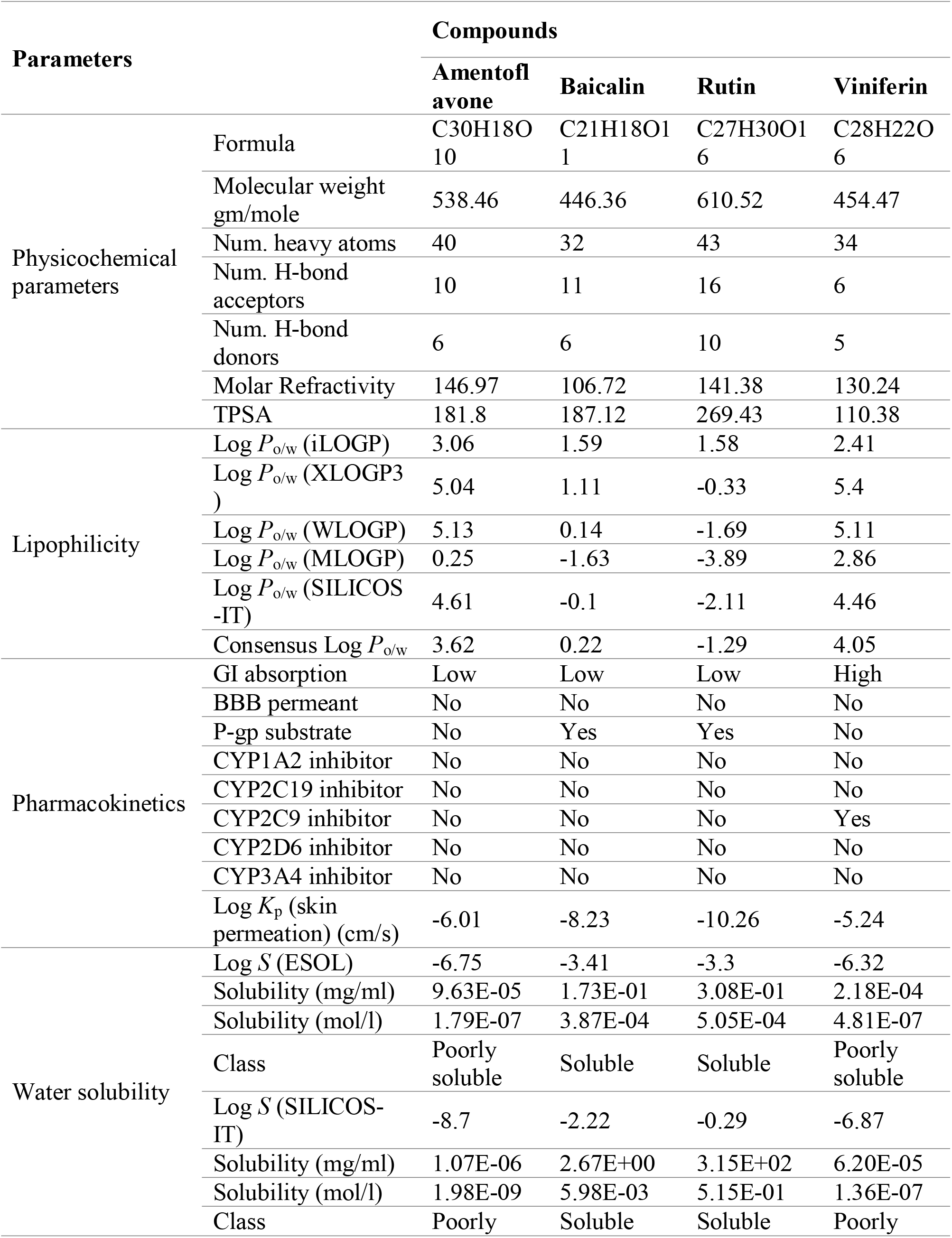
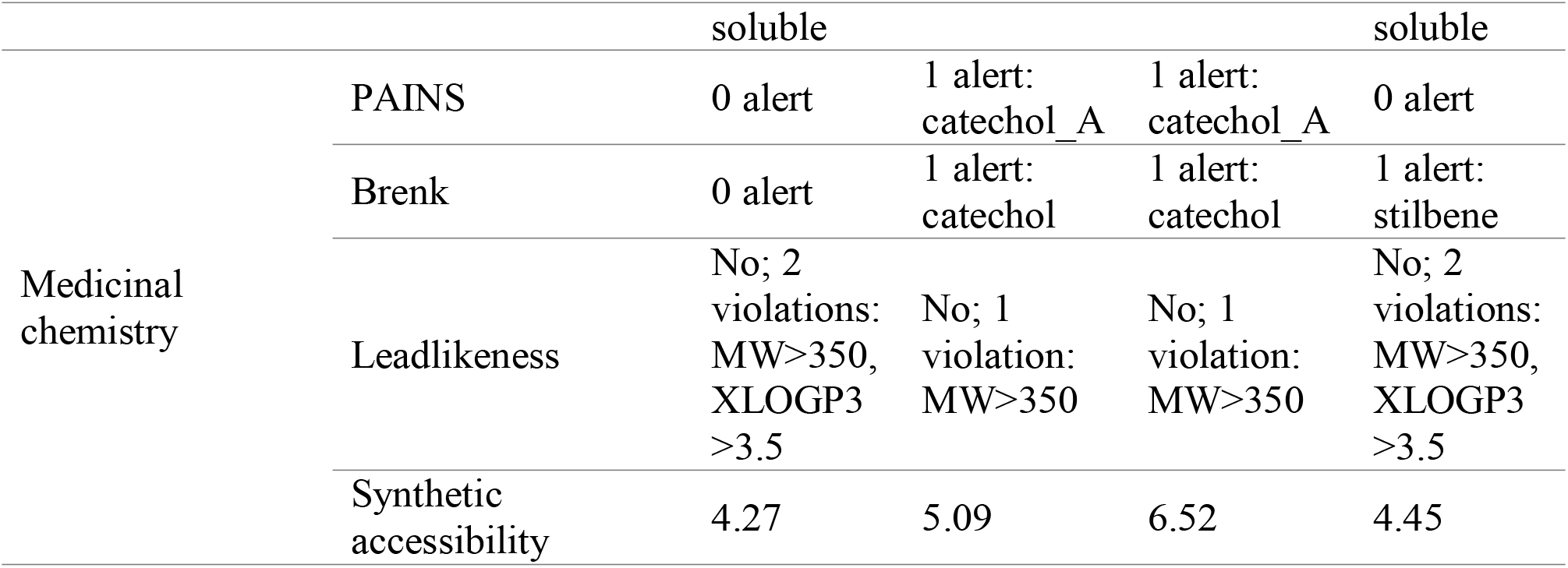
Major metabolites with SwissADME characteristics

### 3.11 Toxicity Assessment

Table-06 displays the toxicity assessments of the top metabolites. The LD50 values for the best drugs were between 2.491 and 2.634.mol/kg, they showed no skin sensitivity and no acute toxicity in rats when administered orally. Minnow All drugs had toxicity values greater to -0.3 log mM, demonstrating that they are not toxic. Once more, all drug tests for hepatotoxicity were negative, demonstrating that the liver’s normal operation won’t be jeopardized by these powerful medications.

**Table 06:** Top metabolites with toxicity prediction

| Toxicity Parameters | Compounds |  |  |  |
| --- | --- | --- | --- | --- |
|  | Amentoflavone | Baicalin | Rutin | Viniferin |
| AMES Toxicity | No | No | No | No |
| Max. Tolerated Dose (log mg/kg/day) | 0.438 | 0.652 | 0.452 | 0.409 |
| hERG I inhibitor | No | No | No | No |
| hERG II inhibitors | Yes | No | Yes | Yes |
| Oral Rat Acute Toxicity, LD50 (mol/kg) | 2.527 | 2.634 | 2.491 | 2.568 |
| Oral Rat Chronic Toxicity,LOAEL (log mg/kg_bw/day) | 3.572 | 4.536 | 3.673 | 3.267 |
| Hepatotoxicity | No | No | No | No |
| Skin Sensitisation | No | No | No | No |
| Minnow Toxicity (log mM) | 2.685 | 5.525 | 7.677 | 1.161 |

## DISCUSSION

*Cryptococcus gattii* is an encapsulated yeast prevalent in tropical and subtropical regions. The lungs are first affected by the infection cryptococcosis, then it rapidly invades the brain and other organs, often with catastrophic consequences. There are very few antifungal medications for C. *gattii* infection, and it has already started to show resistance against one the antifungal drug fluconazole. Therefore, we investigated the proteomes to identify potential drug targets and candidates to combat this pathogenic yeast. Essential proteins are adjudged to be the most ideal antimicrobial therapeutic targets as most of the drugs have a propensity to dock along with essential gene products. The complete proteome of C. *gattii* was retrieved from NCBI. Which is consist of 6561 proteins. We have removed the paralog proteins with 60% match considering that paralogs may share similar pattern with motifs, domains, and active site. Drug that is developed and administrated to bind with target protein of pathogens may cross-react with host homologous proteins (Sarkar et al., 2012). To avoid this, homologous proteins (3800) were excluded and only non-homologous protein (2342) were kept. A potential therapeutic target should be an Essential protein as it possesses crucial quality for the pathogen to exist (Sarkar et al., 2012). A total of 216 protein found essential for C. *gattii* that can cause the death of particular agent. The potential target protein must possess no similar metabolic pathway with the host human to avoid blocking of human’s metabolic pathway which could be fatal. To find out the unique metabolic pathway, it was found that, 117 out of 216 important non-homolog proteins were involved in both metabolic pathways and orthology and only 7 are solely associated with C. *gattii* unique pathway. A determination of the host microbiome’s dissimilarity ensures that those beneficial microbes are protected from unintended blocking. Here, 6 proteins are identified with no similarity with human microflora and only three were found to be very crucial for C. *gattii’*s metabolic cycle and selected as novel drug targets. The selected proteins are situated in plasma membrane, mitochondria and nuclear sites. These proteins play vital functions like protein biogenesis, mitochondrial morphogenesis and mitochondrial inheritance. Moreover, these proteins are also associated with some crucial pathways such as major beta-barrel assembly pathway (D’Souza et al., 2011) making them ideal as drug targets.

Plant metabolites significantly contributes as a predominant molecule to uncover convenient drug contenders (Joseph et al., 2017).So, for C. *gattii* few inhibitory metabolites obtained from plant sources were assessed on the ground of their propensity of joining with certain distinct drug targets. Then the preferred drug targets were docked against chosen plant metabolites, which revealed that 4 molecules Amentoflavone, Baicalin, Rutin, and Viniferin has the lowest global binding energy and the highest affinity for all three selected drug targets. Each of them showed better performance than already resistant drug fluconazole. The structural topologies of the docked protein complex were carefully inspected to comprehend the outward drug hotspot of the targeted unique proteins. The binding configuration of the ligand and related residues, along with their associated sites were assessed (Table 4) (Figure 6, 7 and 8). Improper ADME data is usually associated with the defect of clinical trials over the course of numerous drug development initiatives (Shin et al., 2017). Therefore, it is vital to inspect ADME data to understand the safety and efficacy of a drug candidate. The top four medication candidates displayed no unfavorable effects in the in-silico ADME investigation which might minimize the drug related qualities (Table 05). Among 4 candidates, only amentoflavone and viniferin showed low solubility, while baicalin and rutin had significant solubility. The toxicity prediction displayed that all 4 chosen drug nominees are non-carcinogenic, non-mutagenic, no hepatotoxic and not skin-sensitive. All in all, the toxicity study reveal that predicted novel drugs are totally safe to perform procedures and serve as therapeutic medication to treat C. *gattii*.

## Conclusion

Cryptococcosis infection caused by *Cryptococcus gattii* might result in fatal consequences. It is already showing resistance against existing drugs. In order to develop new antifungal medicines, it is necessary to investigate potential pharmacological targets that can help slow down the fast growth of antifungal resistance among *Cryptococcus gattii*. Through this study, three proteins of this fungus were identified as potential prospective therapeutic targets. The development of novel antifungal medicines that would block these prospective drug targets might effectively prevent the spread of *Cryptococcus gattii*. However, these putative drug targets still need to be further examined and experimentally confirmed.

## Data Availability

The data used to support the findings of this study are available from the corresponding author upon request.

## Authors’ Contributions

Tanjin Barketullah Robin, Nurul Amin Rani, Nadim Ahmed, Anindita Ash Prome and Md. Nazmul Islam Bappy contributed equally to this work under Foeaz Ahmed supervision.

