## Supplementary File S4 for "In Silico Proteomics Approach Towards the Identification of Potential Novel Drug Targets Against *Cryptococcus gattii*"

Supplementary File S4: Unique metabolic pathways of C.*gattii*.

| **Entry** | | **Name** | **Description** | **Object** | **Legend** | |
| --- | --- | --- | --- | --- | --- | --- |
| [cgi00261](https://www.genome.jp/dbget-bin/www_bget?pathway:cgi00261) | | Monobactam biosynthesis | Monobactams are beta-lactam antibiotics containing a monocyclic beta-lactam nucleus, which is struct... | C01179 (3-(4-Hydroxyphenyl)pyruvate) C03198 ((S)-4-Hydroxymandelate) C03590 (4-Hydroxyphenylglyoxyla... | MONOBACTAM BIOSYNTHESIS Phenylalanine, tyrosine and tryptophan biosynthesis 4-Hydroxy-phenylpyruvat... | |
| [cgi00300](https://www.genome.jp/dbget-bin/www_bget?pathway:cgi00300) | | Lysine biosynthesis |  | C19889 (LysW-gamma-L-lysine) C19888 (LysW-gamma-L-alpha-aminoadipate 6-semialdehyde) C19887 (LysW-ga... | 2.6.1.83 Tropane, piperidine and pyridine alkaloid biosynthesis LysW-γ-L-lysine LysW-γ-L-α-aminoa... | |
| [cgi00332](https://www.genome.jp/dbget-bin/www_bget?pathway:cgi00332) | | Carbapenem biosynthesis | Carbapenems are broad-spectrum beta-lactam antibiotics, which are often considered as the antibiotic... | C00025 (L-Glutamate) C03287 (L-Glutamyl 5-phosphate) C01165 (L-Glutamate 5-semialdehyde) C03912 ((S)... | CARBAPENEM BIOSYNTHESIS Arginine and proline metabolism L-Glutamate 2.7.2.11 L-Glutamyl-P 1.2.1.41 ... | |
| [cgi00460](https://www.genome.jp/dbget-bin/www_bget?pathway:cgi00460) | | Cyanoamino acid metabolism |  | C05670 (3-Aminopropiononitrile) C01401 (Alanine) C00302 (Glutamate) C05714 (alpha-Aminopropiononitri... | 3-Aminopropiono-nitrile D-Amino acid metabolism 3.5.5.1 3.5.5.1 CYANOAMINO ACID METABOLISM β-Alan... | |
| [cgi00660](https://www.genome.jp/dbget-bin/www_bget?pathway:cgi00660) | | C5-Branched dibasic acid metabolism |  | C00810 ((R)-Acetoin) C06010 ((S)-2-Acetolactate) C01011 ((3S)-Citramalyl-CoA) C00531 (Itaconyl-CoA) ... | | Nicotinate and nicotinamide metabolism Valine, leucine and isoleucine biosynthesis 2.3.3.11 Alanine,... |
| [cgi00680](https://www.genome.jp/dbget-bin/www_bget?pathway:cgi00680) | Methane metabolism | | Methane is metabolized principally by methanotrophs and methanogens in the global carbon cycle. Meth... | C01438 (Methane) C00132 (Methanol) C00067 (Formaldehyde) C00058 (Formate) C00237 (CO) C04330 (5,10-M... | | METHANE METABOLISM Methane 1.14.13.25 1.14.18.3 Methanol Formaldehyde 1.1.3.13 1.1.2.7 1.1.1.244 1.... |

| [cgi00791](https://www.genome.jp/dbget-bin/www_bget?pathway:cgi00791) | Atrazine degradation |  | C01566 (Cyanamide) C08737 (Melamine) C08733 (Ammeline) C08734 (Ammelide) C14149 (N-Cyclopropylammeli... | 4.2.1.69 Cyanamide Melamine Ammeline Ammelide N-Cyclopropyl-ammelide N-Cyclopropyl-ammeline 3.8.1.8 ... |
| --- | --- | --- | --- | --- |
| [cgi00909](https://www.genome.jp/dbget-bin/www_bget?pathway:cgi00909) | Sesquiterpenoid and triterpenoid biosynthesis | Sesquiterpenoids (C15 terpenoids) are a group of terpenoids consisting of three isoprene units. They... | C17954 (Albaflavenone) C00448 (trans,trans-Farnesyl diphosphate) C09704 (Nerolidol) C09665 (alpha-Fa... | Albaflavenone Terpenoid backbone biosynthesis Farnesyl-PP (S,E)-Nerolidol (E,E)-α-Farnesene (E,E)-... |
| [cgi00999](https://www.genome.jp/dbget-bin/www_bget?pathway:cgi00999) | Biosynthesis of various plant secondary metabolites |  | C22445 (8-Hydroxybergapten) C09315 (Umbelliferone) C18083 (Demethylsuberosin) C09276 (Marmesin) C007... | 2.5.1.139 1.14.14.66 1.14.14.141 1.14.13.- 2.1.1.69 2.5.1.139 1.14.14.148 2.1.1.70 1.14.11.60 8-Hydr... |
| [cgi01110](https://www.genome.jp/dbget-bin/www_bget?pathway:cgi01110) | Biosynthesis of secondary metabolites |  | C20799 (Grixazone B) C22416 (Cremeomycin) C22425 (3-Amino-2-hydroxy-4-methoxybenzoate) C22455 (3-Ami... | Biosynthesis of various plant secondary metabolites α-Linolenic acid metabolism Type I polyketide s... |
| [cgi01232](https://www.genome.jp/dbget-bin/www_bget?pathway:cgi01232) | Nucleotide metabolism |  | C00130 (IMP) C00104 (IDP) C00081 (ITP) C00020 (AMP) C00008 (ADP) C00002 (ATP) C00294 (Inosine) C0021... | IMP IDP ITP AMP ADP ATP Inosine Adenosine Adenine Hypoxanthine Guanine GDP GMP Guanosine GTP Xanthin... |
| [cgi03250](https://www.genome.jp/dbget-bin/www_bget?pathway:cgi03250) | Viral life cycle - HIV-1 |  | CGB_C3340C CGB_C1480C, CGB_D6570W CGB_G6440W CGB_N0130W CGB_A8280C CGB_C5090C CGB_N0130W CGB_A8280C ... | Binding Preintegration complex (PIC) reverse transcriptase matrix integrase Fusion Reverse trans... |
| [cgi04011](https://www.genome.jp/dbget-bin/www_bget?pathway:cgi04011) | MAPK signaling pathway - yeast | The S. cerevisiae genome encodes multiple MAP kinase orthologs. One (Fus3) mediates cellular respons... | CGB_I1200C CGB_E0290W, CGB_E0450W CGB_C3420C CGB_I1040C CGB_F6480W CGB_F6460W, CGB_H0200C CGB_L1550C... | Tec1 Ste12 Dig1,2 Kss1 Ste7 Ste11 Ste20 Cdc42 Sho1 Mlp1 Slt2 Tus1 Rho1 Wsc1,2,3 Pkc1 Bck1 Mkk1,2 Rlm... |
| [cgi04111](https://www.genome.jp/dbget-bin/www_bget?pathway:cgi04111) | Cell cycle - yeast | Mitotic cell cycle progression is accomplished through a reproducible sequence of events, DNA replic... | C00575 (3',5'-Cyclic AMP) C00009 (Orthophosphate) CGB_A0440W, CGB_A1490W, CGB_B1120C, CGB_F4520C, CG... | MCM ORC APC/C SCF SCF cAMP Phosphate Ubiquitin mediated proteolysis Mcm7 Mcm6 Mcm5 Mcm4 Mcm3 Mcm2 Or... |
| [cgi04113](https://www.genome.jp/dbget-bin/www_bget?pathway:cgi04113) | Meiosis - yeast | During meiosis, a single round of DNA replication is followed by two rounds of chromosome segregatio... | C00031 (D-Glucose) C00031 (D-Glucose) C00288 (HCO3-) C00575 (3',5'-Cyclic AMP) CGB_F4520C CGB_A1490W... | Mcm7 Mcm6 Mcm5 Mcm4 Mcm3 Mcm2 Orc6 Orc5 Orc4 Orc3 Orc2 Orc1 MEIOSIS - YEAST MCM (Mini-Chromosome M... |
| [cgi04138](https://www.genome.jp/dbget-bin/www_bget?pathway:cgi04138) | Autophagy - yeast | Autophagy is a non-selective and bulk intracellular degradation system of eukaryotic cells and is hi... | C04549 (1-Phosphatidyl-1D-myo-inositol 3-phosphate) C01194 (1-Phosphatidyl-D-myo-inositol) C00350 (P... | PI3P ATG6 VPS15 ATG17 ATG1 ATG9 ATG13 ATG14 VPS34 Snf1 AUTOPHAGY - YEAST Nutrient starvation ATG31... |
| [cgi04139](https://www.genome.jp/dbget-bin/www_bget?pathway:cgi04139) | Mitophagy - yeast | Mitophagy, which refers to the selective elimination of impaired or excessive mitochondria, is consi... | D00753 (Sirolimus (JAN/USAN/INN)) CGB_I2500W CGB_E2440W CGB_H1030C CGB_E3150C, CGB_F2520W, CGB_G6520... | MITOPHAGY - YEAST WSC1 PKC1 BCK1 MKK1/2 SLT2 ATG32 Excess ROS Oxidative stress DNA SLN1 CK2 HOG1 S... |
