## Supplementary File S5 for "In Silico Proteomics Approach Towards the Identification of Potential Novel Drug Targets Against *Cryptococcus gattii*"

Supplementary File S5: List of plant metabolites with antifungal activity.

| SL No. | Metabolites | Class | PubChem CID | Source | Biological Effects | References |
| --- | --- | --- | --- | --- | --- | --- |
| 1. | Ajoene | Organosulfur compound | 5386591 | *Allium sativum* | Anti-viral, Anti-thrombotic, anti-tumoral, anti-fungal, and anti-parasitic effects | (Weber et al., 1992) |
| 2. | Thymol | Monoterpene phenol | 6989 | Thymus vulgaris | Anti-cancer, anti-microbial, anti-viral, anti-oxidant, immunomodulators, anti-inflammatory, anti-fungal and anti-spasmodic | (Ranjbar, Ramezanian, Shekarforoush, Niakousari, & Eshghi, 2022) |
| 3. | Carvacrol | Monoterpene phenol | 10364 | Origanum vulgare | Anti-oxidant, anti-microbial, anti-viral, anti-hypertensive, immunomodulatory and anti-cancer | (Oliveira Lima et al., 2013) |
| 4. | Caffeic acid | Phenolic compound | 689043 | Eucalyptus globulus | Anti-oxidant, reducing, anti-fungal, anti-ACE(angiotensin-converting enzyme ) anti-inflammatory and chelating activity | (Vio-Michaelis, Apablaza-Hidalgo, Gómez, Peña-Vera, & Montenegro, 2012) |
| 5. | Linalool | Acyclic monoterpenoid | 6549 | *Ocimum basilicum* | Anti-cancer, anti-microbial, neuroprotective, anxiolytic, antidepressant, anti-stress, hepatoprotective, renal protective, and lung protective , anti-inflammatory activity | Meena et al.(2013);  An et al. (2021). |
| 6. | Jatrorrhizine | Alkaloids | 72323 | *Mahonia aquifol* | Anticancer, anti-fungal, anti-microbial, detoxification, bactericidal, hypoglycemic, hypolipidemic and antiparasitic properties | Jung et al. (2006); Li et al. (2014); Jiang et al. (2011); Wang et al. (2018); Wang et al. (2019) |
| 7. | Protopine | Alkaloids | 4970 | Fumaria vaillantii | Anti-inflammatory, anti-platelet aggregation, anti-cancer, analgesic, vasodilatory, anti-cholinesterase, anti-addictive, anticonvulsant, anti-pathogenic, anti-oxidant, hepatoprotectiv, neuroprotective, and cytotoxic and anti-proliferative activities. | (Qing et al., 2017) |
| 8. | Viniferin | Stilbenoid | 5315232 | Dryobalanops aromatica. | Anti-viral, anti-fungal, anti-inflammatory, anti-bacterial, anti-cancer and anti-oxidant activity | (Langcake, 1981) |
| 9. | abscisic acid | isoprenoid | 5375199 | *Lygodium japonicum* | Antidiabetic effect, antifungal effect | (Khedr, Massarotti, & Mohamed, 2018) |
| 10. | Rutin | flavonoid | 5280805 | *Styphnolobium japonicum* | antioxidant and anti-inflammatory effects | (Oliveira, Carraro, Auler, & Khalil, 2016) |
| 11. | Gallic acid | phenolic | 370 | *Myriophyllum spicatum* | antioxidant, anti-inflammatory, and antineoplastic propertie | (Li et al., 2017) |
| 12. | Eugenol | allylbenzene | 3314 | *Eugenia caryophyllata* | antimicrobial, anti-inflammatory, analgesic, neuroprotective, antidiabetic, Anti-fungal | (Hassanpour, Shams-Ghahfarokhi, & Razzaghi-Abyaneh, 2020)  (Wang, Zhang, Chen, Fan, & Shi, 2010) |
| 13. | Quercetin | Flavanol | 5280343 | Aloe vera (L.) Burm.f. | Antiviral activity, Anti-inflammatory, Antioxidant | (Oliveira et al., 2016) |
| 14. | Amentoflavone | Polyphenolic | 5281600 | Torreya nucifera | Antiviral, Antisenescence, Antioxidation | (Jung et al., 2006)  (Hwang, Lee, Jin, Woo, & Lee, 2012) |
| 15. | Baicalin | Flavonoid | 64982 | Scutellaria. baicallensis | Antiviral activity, anti-inflammatory activity | (Serpa et al., 2012) |
| 16. | Betulinic acid | triterpenoid | 64971 | Betula pubescens | Anti-viral, Antimalarial, Anti-cancer activity | (Vaishnav & Demain, 2011) |
| 17. | Resveratrol | Stilbenoid | 445154 | *Vitis vinifera* | Anti-tumor, anti-aging, anti-inflammatory, anti-oxidant, anti-radiation, immune regulation, cardiovascular protection,  anti-fungal properties etc. | Meena et al.(2013); CAO, et al. (2021) |
| 18. | 3-methoxysampangine | Alkaloids | 122683 | *Cleistopholis patens* | Anti-malarial, anti-fungal, and cytotoxic potency | Meena et al.(2013); Liu et al. (1990) |
| 19. | Usnic acid | Benzofurans | 5646 | Parmelia perlata (Huds.) Ach | Anti-inflammatory, analgesic, healing, anti-oxidant, anti-microbial, anti-protozoal, anti-viral, larvicidal and UV protection | Khare(2007); Araújo et al. (2015). |
| 20. | 4-Methyl-7-hydroxycoumarin | hydroxycoumarin | 5280567 | *Dipteryx odorata* | Antifungal, Antioxidant, Anti-cancer, anti-microbial, Anti-tubercular | (Sarkanj, Molnar, Čačič, & Gille, 2013) |
| 21. | Eucalyptol | monoterpenoid | 2758 | *Eucalyptus spp* | Antifungal, Anti-inflammatory, Antioxidant, Anticancer, antibacterial, Anti-malarial | (Sarkanj et al., 2013) |
