## Supplementary File S6 for "In Silico Proteomics Approach Towards the Identification of Potential Novel Drug Targets Against *Cryptococcus gattii*"

Supplementary File S6: Molecular docking of protein-ligand complex.

| **SI No** | **Protein Name** | **Name of Metabolites** | **Docking Energy** | **RMSD Value** |
| --- | --- | --- | --- | --- |
| 1. | XP 003194316.1 | 3-methoxysampangine | -144.57 | 113.93 |
| 2. |  | 4-Methyl-7-hydroxycoumarin | -115.16 | 120.15 |
| 3. |  | abscisic acid | -142.31 | 117.03 |
| 4. |  | Ajoene | -83.23 | 118.76 |
| 5. |  | Amentoflavone | -237.02 | 115.76 |
| 6. |  | Baicalin | -200.13 | 117.08 |
| 7. |  | Betulinic acid | -159.37 | 123.59 |
| 8. |  | Caffeic acid | -127.85 | 120.60 |
| 9. |  | Carvacrol | -102.93 | 120.08 |
| 10. |  | Eucalyptol | -93.45 | 114.59 |
| 11. |  | Eugenol | -102.27 | 113.88 |
| 12. |  | **Fluconazole** | **-153.40** | **114.45** |
| 13. |  | Gallic acid | -111.90 | 120.46 |
| 14. |  | Jatrorrhizine | -162.95 | 116.04 |
| 15. |  | Linalool | -90.96 | 120.63 |
| 16. |  | Protopine | -177.09 | 115.66 |
| 17. |  | Quercetin | -175.13 | 120.62 |
| 18. |  | Resveratrol | -142.31 | 117.42 |
| 19. |  | Rutin | -234.36 | 109.61 |
| 20. |  | Thymol | -103.98 | 120.25 |
| 21. |  | Usnic acid | -168.99 | 114.82 |
| 22. |  | Viniferin | -187.22 | 109.38 |
| 23. | XP 003197297.1 | 3-methoxysampangine | -125.69 | 101.67 |
| 24. |  | 4-Methyl-7-hydroxycoumarin | -95.94 | 110.87 |
| 25. |  | abscisic acid | -120.53 | 97.64 |
| 26. |  | Ajoene | -68.86 | 102.52 |
| 27. |  | Amentoflavone | -197.08 | 107.64 |
| 28. |  | Baicalin | -201.20 | 101.42 |
| 29. |  | Betulinic acid | -151.04 | 97.47 |
| 30. |  | Caffeic acid | -98.28 | 110.98 |
| 31. |  | Carvacrol | -77.66 | 97.67 |
| 32. |  | Eucalyptol | -78.52 | 111.51 |
| 33. |  | Eugenol | -85.15 | 110.48 |
| 34. |  | **Fluconazole** | **-122.25** | **98.15** |
| 35. |  | Gallic acid | -97.94 | 111.28 |
| 36. |  | Jatrorrhizine | -150.41 | 102.46 |
| 37. |  | Linalool | -76.63 | 98.39 |
| 38. |  | Protopine | -150.24 | 104.25 |
| 39. |  | Quercetin | -151.31 | 99.83 |
| 40. |  | Resveratrol | -113.65 | 98.01 |
| 41. |  | Rutin | -192.27 | 107.51 |
| 42. |  | Thymol | -79.40 | 97.68 |
| 43. |  | Usnic acid | -144.06 | 104.76 |
| 44. |  | Viniferin | -171.08 | 100.60 |
| 45. | XP 003197520.1 | 3-methoxysampangine | -160.53 | 129.01 |
| 46. |  | 4-Methyl-7-hydroxycoumarin | -125.52 | 130.00 |
| 47. |  | abscisic acid | -153.57 | 125.21 |
| 48. |  | Ajoene | -84.66 | 128.88 |
| 49. |  | Amentoflavone | -253.13 | 126.42 |
| 50. |  | Baicalin | -239.98 | 127.56 |
| 51. |  | Betulinic acid | -197.54 | 126.54 |
| 52. |  | Caffeic acid | -120.46 | 130.07 |
| 53. |  | Carvacrol | -101.11 | 126.61 |
| 54. |  | Eucalyptol | -92.51 | 124.23 |
| 55. |  | Eugenol | -108.75 | 127.27 |
| 56. |  | **Fluconazole** | **-158.78** | **124.17** |
| 57. |  | Gallic acid | -120.92 | 130.31 |
| 58. |  | Jatrorrhizine | -189.84 | 127.20 |
| 59. |  | Linalool | -95.01 | 125.62 |
| 60. |  | Protopine | -189.95 | 123.77 |
| 61. |  | Quercetin | -197.98 | 127.07 |
| 62. |  | Resveratrol | -152.01 | 127.74 |
| 63. |  | Rutin | -236.93 | 123.43 |
| 64. |  | Thymol | -97.49 | 128.82 |
| 65. |  | Usnic acid | -199.99 | 128.18 |
| 66. |  | Viniferin | -233.25 | 124.98 |
